# Resolving how genetic perturbations act through gene regulatory paths

**DOI:** 10.64898/2026.11.02.763749

**Authors:** Lixin Ren, Zhana Duren

## Abstract

Perturb-seq measures transcriptional responses to thousands of genetic perturbations, but each gene’s response mixes the contributions of many regulatory paths, so how a perturbation acts cannot be resolved from expression alone, however large the model. Here we present REGA-perturb, which supplies this missing information rather than adding model complexity: reference epigenomes define a scaffold of possible transcription factor (TF)–regulatory element–gene paths, and perturbation responses, pooled through shared latent programs, determine how each path is used, requiring only RNA measurements. Across three CRISPR interference screens totaling 2.4 million cells, REGA-perturb inferred TF–target relationships that agreed with ChIP-seq binding with a gain over random more than four times that of the best existing method, separated candidate direct from indirect targets, and predicted targets of TFs withheld from training. It also inferred perturbation-induced changes in TF activity not reflected in TF transcript levels, and resolved perturbation-specific regulatory rewiring in which opposing regulatory changes cancel at the expression level.

## Introduction

Genetic perturbations provide a direct means of investigating gene function and the organization of regulatory relationships. Perturb-seq couples pooled CRISPR perturbations with single-cell RNA sequencing, enabling the transcriptional consequences of thousands of perturbations to be measured in a single experiment^1–3^, and genome-scale screens comprising millions of cells are now available as public resources^3^. The goal of such screens is to understand how perturbations act. However, the expression change of each gene is the net outcome of multiple regulatory inputs, combining direct effects with secondary effects mediated by other regulators^4,5^. The response therefore reflects regulatory mechanism but mixes the contributions of many regulatory paths, so the response alone cannot reveal which paths produced it. Even for the most direct perturbation, knockdown of a transcription factor (TF), genes bound by the TF often show little or no response^5–7^, and the magnitude of expression changes is largely unrelated to where the TF binds, as we confirm across hundreds of TF perturbations in this study. Because many different combinations of regulatory paths can produce the same expression response, this is a limitation of the information in the data rather than of model capacity, and it cannot be overcome simply by building larger models. Yet computational analysis of Perturb-seq data has largely measured progress by how accurately models predict perturbation responses, including responses to unseen perturbations, perturbation combinations and new cellular contexts^8–12^, a criterion that can be met without resolving mechanism. Recent benchmarks have further found that increasingly complex prediction models often do not outperform simple linear baselines^13^.

As a result, neither expression changes nor predictive models reveal which TFs regulate which genes, or through which regulatory elements (REs), and without such a map several questions central to interpreting a perturbation screen cannot be answered. First, when a regulator gene is perturbed, which of the responding genes does it regulate directly, and which respond only secondarily through other regulators? Second, how does a perturbation alter the activity of TFs? Most genes perturbed in a screen are not TFs, yet they can still change TF activity, for example by disrupting cofactors, chromatin regulators or signaling pathways that control TF function, or by triggering stress responses that activate specific TFs^14,15^. Because such changes often act at the protein level, TF activity need not be reflected in TF transcript levels^16^. Third, can the regulatory inputs to a gene change without changing its expression? If a perturbation strengthens some inputs to a gene while weakening others, their effects may cancel, leaving expression stable even though the gene’s regulation has been substantially reorganized. Such compensation is a recognized feature of regulatory systems, arising for example from feedback loops and from redundant regulators that offset the loss of one another^17,18^. Yet because it leaves no trace in expression, it is invisible to differential expression analysis, which would classify these genes as unaffected. A regulatory map that answers these questions would also be most useful if it reached beyond the perturbations actually performed. Most TFs expressed in a cell type are never perturbed in a given screen, and a map that captures shared regulatory structure, rather than the idiosyncrasies of individual perturbations, should anticipate their effects as well.

Existing approaches that aim to model regulatory mechanism address this problem only in part. Latent program analyses identify coordinated transcriptional responses to perturbations^19^ but do not link these programs to the TFs and REs that produce them. Gene regulatory network (GRN) inference methods, including SCENIC, CellOracle and GRNBoost2^20–22^, do model regulatory relationships, but they infer them largely from covariation in expression, in which the contributions of different regulatory paths remain confounded. They show limited agreement with TF binding in benchmarking studies^23,24^ and yield static networks that do not describe how individual relationships change across perturbations. Most also do not exploit perturbations as interventions on a shared regulatory structure. Methods that additionally use single-cell chromatin accessibility, such as LINGER and SCENIC+^25,26^, cannot learn from Perturb-seq screens, which measure only RNA, and when applied to unperturbed cells, they share the same static limitation.

Resolving these contributions requires information that expression data alone cannot provide: which regulatory paths can exist. This information is available from a complementary source. Reference epigenomic catalogs from large-scale projects such as ENCODE^27^, combined with TF motifs and RE– gene proximity, specify candidate regulatory paths linking TFs to target genes through REs^28^, providing a scaffold of possible regulation. These catalogs define only potential relationships, however, not which of them are used in a given cell or how their use changes under perturbation. Perturb-seq provides this complementary information: the consequences of thousands of interventions, without the paths responsible. Integrating the two through latent regulatory programs shared across perturbations could restrict inference to biologically plausible paths, pool information across thousands of interventions, and allow the contribution of each path to vary from one perturbation to another. Because the resulting paths make explicit mechanistic claims, they can also be evaluated directly against independent measurements of regulation, such as TF binding and chromatin contacts, rather than by prediction accuracy alone.

Here we present REGA-perturb (Reference REgulatory Path-Guided Analysis), which supplies this missing information rather than adding model complexity. REGA-perturb represents each perturbation as an activity profile over shared latent regulatory programs and propagates these activities to downstream genes through TF motifs and REs defined by a cell-type-matched reference epigenome, learning from perturbation data how strongly each path is used. It requires only RNA measurements and scales to millions of cells, so it can be applied directly to existing Perturb-seq screens. Across three CRISPRi screens totaling 2.4 million cells, REGA-perturb inferred TF–target relationships that agreed with ChIP-seq binding substantially better than existing methods, and predicted regulons responsive to TFs withheld from training. It also addressed each of the three questions above: it separated candidate direct from indirect targets of perturbed TFs, inferred how perturbations alter TF activity in ways not reflected in TF transcript levels, and resolved perturbation-specific regulatory rewiring, including opposing changes in regulatory inputs that cancel at the expression level.

## Results

### Overview of REGA-perturb

We developed REGA-perturb, a Reference REgulatory Path-Guided Analysis framework for tracing single-cell perturbation responses through gene regulatory paths. Rather than directly modeling perturbation–gene relationships, REGA-perturb models the propagation of perturbation signals through a hierarchical regulatory architecture, linking perturbation targets to downstream transcriptional responses via latent regulatory programs, transcription factor (TF) motifs, regulatory elements (REs) and target genes (Fig. 1a).

**Figure 1.**
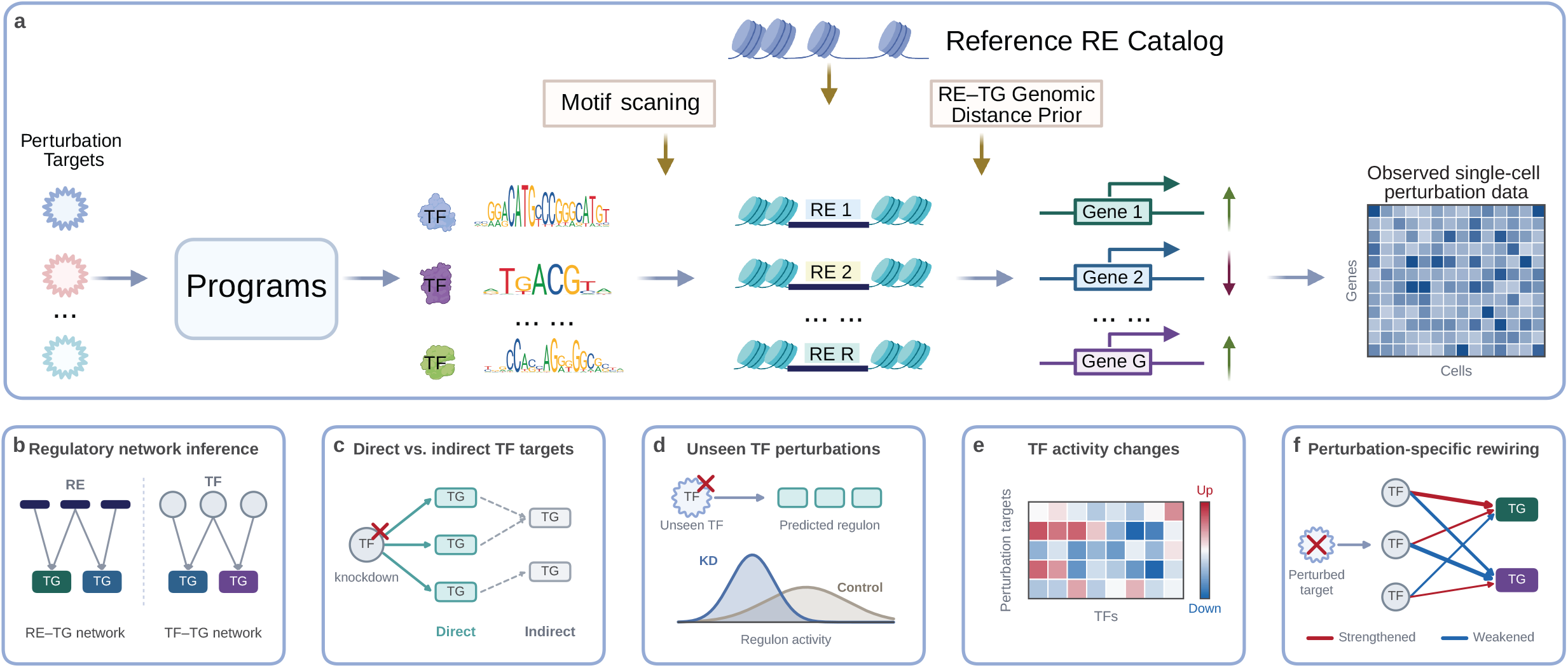
Overview of the REGA-perturb framework. **a**, Schematic of the REGA-perturb model, which traces how the effect of a genetic perturbation propagates through a hierarchical regulatory chain to shape single-cell gene expression. Information flows from perturbation targets to latent regulatory programs, then through transcription factor (TF) motifs and regulatory elements (REs) to downstream target genes (TGs), together reconstructing the observed gene-by-cell expression matrix of a Perturb-seq experiment. The TF motif → RE → TG part of this chain (the reference regulatory path) is constrained by motif scanning of a reference RE catalog and by an RE–TG prior based on genomic distance. **b–f**, Downstream analyses supported by REGA-perturb. **b**, Inference of RE–TG and TF–TG regulatory networks. **c**, Identification of candidate direct and indirect targets of perturbed TFs. **d**, Identification of regulons responsive to unseen TF perturbations, evaluated by comparing regulon activities between knockdown and control cells. **e**, Characterization of perturbation-associated changes in TF activity. **f**, Analysis of perturbation-specific regulatory rewiring, highlighting increases and decreases in inferred TF–TG regulatory influence.

A central feature of REGA-perturb is the incorporation of a reference regulatory path connecting TF motifs to target genes through REs. This path is constructed from a cell-type-matched reference RE catalog, with motif occurrences defining candidate TF motif–RE associations and genomic proximity defining candidate RE–gene associations. The reference regulatory path provides a structural prior that constrains the regulatory connections considered by the model while allowing their strengths to be refined using Perturb-seq data. By integrating prior regulatory knowledge with perturbation-associated transcriptional profiles, REGA-perturb learns regulatory relationships within a biologically informed framework.

REGA-perturb represents each perturbation target by an activity profile across a shared set of latent regulatory programs. By jointly learning program activities and regulatory weights, REGA-perturb reconstructs observed transcriptional profiles and provides an interpretable decomposition of perturbation-associated regulatory effects. This representation enables the contributions of individual perturbations to downstream gene expression to be traced through specific regulatory programs, TF motifs and REs. Although the regulatory architecture is shared across perturbations, differences in program activity profiles allow the inferred contributions of individual regulatory paths to vary across perturbations.

REGA-perturb is designed to scale to large Perturb-seq datasets and was applied to CRISPR interference (CRISPRi) screens comprising millions of single cells and thousands of perturbation targets. By capturing perturbation-associated regulatory relationships, REGA-perturb supports downstream analyses including gene regulatory network inference (Fig. 1b), identification of candidate direct and indirect targets of perturbed TFs (Fig. 1c), prediction of regulons responsive to unseen TF perturbations (Fig. 1d), characterization of perturbation-induced changes in TF activity (Fig. 1e), and investigation of perturbation-specific regulatory rewiring (Fig. 1f).

### REGA-perturb improves GRN inference from single-cell perturbation data

To evaluate REGA-perturb, we applied it to three large-scale Perturb-seq datasets from two well-studied human cell lines, each comprising more than 2,000 perturbation targets. Two were generated in myelogenous leukemia lymphoblasts (K562): a genome-wide screen (K562 GWPS; ~1.97 million cells and 9,866 perturbation targets) and an essential-gene screen (K562 essential; ~302,000 cells and 2,057 perturbation targets)^3^. The third was generated in liver hepatocellular carcinoma cells (HepG2; ~142,000 cells and 2,393 perturbation targets)^29^. For each dataset, peaks from ENCODE bulk ATAC-seq data of the matching cell line served as the reference RE catalog (see Data Availability). To assess whether REGA-perturb prioritizes functional REs, we compared the REs with the highest inferred regulatory importance scores with ENCODE ChIP-seq peaks^27^ for the active promoter mark H3K4me3^30^ and the active enhancer mark H3K27ac^31^ in the same cell lines. The top-ranked REs were significantly enriched for both marks in all three datasets, with average fold enrichments of 2.36 for H3K27ac and 3.24 for H3K4me3 (Fig. 2a). They were also more evolutionarily conserved: across thresholds from the top and bottom 10% to 50%, REs with high importance had significantly higher sequence conservation scores than REs with low importance in all three datasets (Fig. 2b; Supplementary Fig. 1a,b). These results indicate that the REs to which REGA-perturb assigns high importance are functionally active and evolutionarily conserved, even though neither histone-mark nor conservation data were provided to the model.

**Figure 2.**
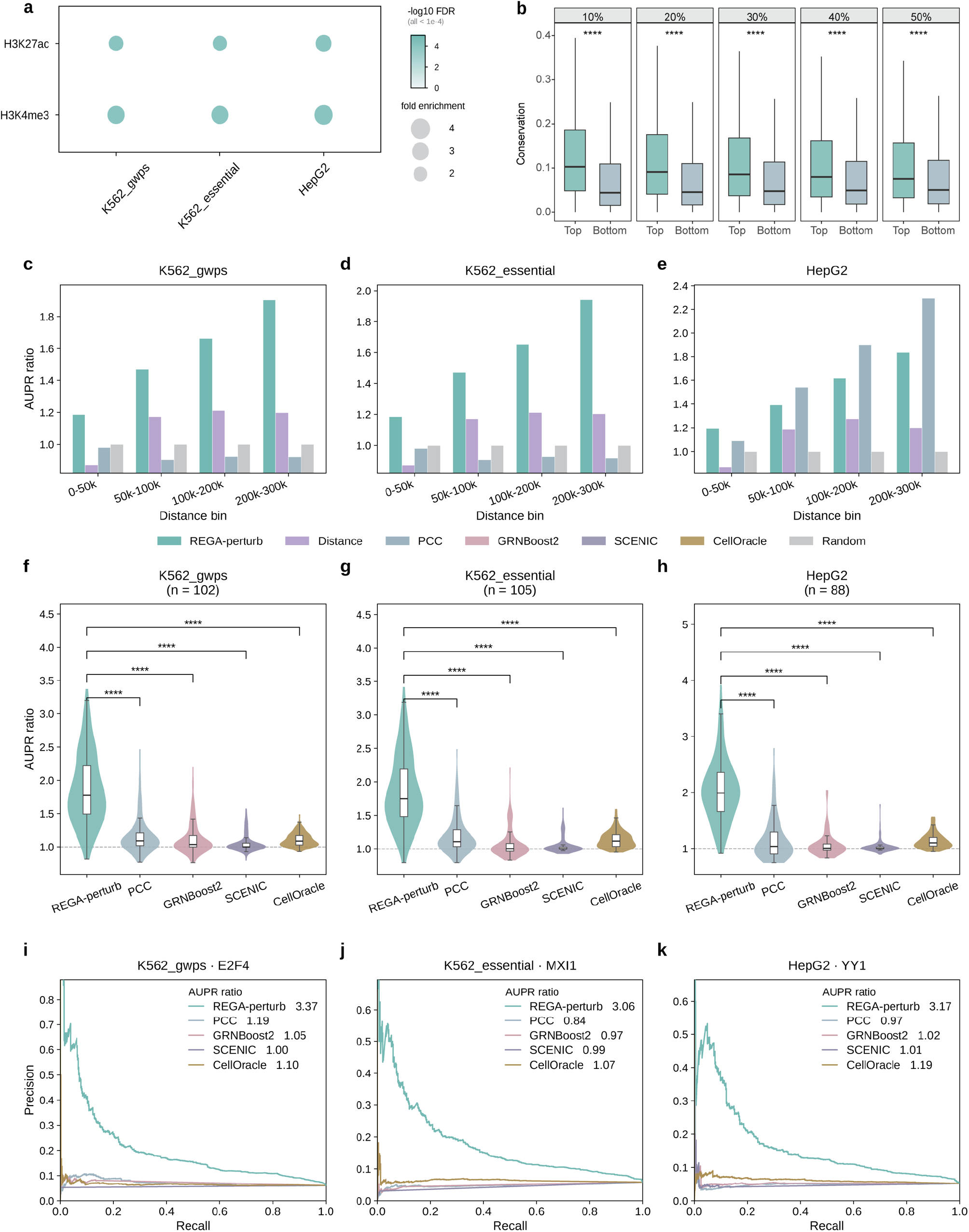
REGA-perturb improves gene regulatory network inference from single-cell perturbation data. **a**, Bubble plot showing the enrichment of the top 2,000 regulatory elements (REs) ranked by REGA-perturb importance in H3K27ac and H3K4me3 ChIP-seq peaks across the K562 genome-wide Perturb-seq screen (K562 GWPS), K562 essential-gene screen (K562 essential) and HepG2 dataset. Bubble size represents fold enrichment relative to 10,000 randomly sampled RE sets of the same size, and color indicates −log_10_(FDR) (one-sided permutation test with Benjamini–Hochberg correction; FDR ≤ 1.0 × 10^−4^ for all comparisons). Fold enrichments for H3K27ac and H3K4me3 were 2.18 and 3.03 in K562 GWPS, 2.18 and 3.11 in K562 essential, and 2.71 and 3.58 in HepG2, respectively. **b**, Boxplots comparing sequence conservation scores (100-way phastCons) between REs with high (Top) and low (Bottom) REGA-perturb importance in K562 GWPS across percentile thresholds of 10%, 20%, 30%, 40% and 50% (two-sided Welch’s t-test; P = 5.4 × 10^−92^, 2.2 × 10^−105^, 1.1 × 10^−115^, 2.7 × 10^−105^ and 1.5 × 10^−91^, respectively; ****P < 0.0001). Boxes indicate the median and interquartile range (IQR), and whiskers extend to the most extreme observations within 1.5 × IQR of the quartiles. **c–e**, Bar charts showing area under the precision–recall curve (AUPR) ratios for predicting RE–target gene (RE–TG) interactions across four genomic distance bins (0–50, 50–100, 100–200 and 200–300 kb from the transcription start site), using RNA polymerase II ChIA-PET interactions as the benchmark reference. Results are shown for K562 GWPS (**c**), K562 essential (**d**) and HepG2 (**e**). REGA-perturb was compared with Distance, PCC (Pearson correlation between RE accessibility and gene expression in independent paired single-cell multiome data) and Random baselines. Each bar represents a single AUPR ratio calculated across all candidate RE–TG pairs within the corresponding distance bin. **f–h**, Violin plots comparing AUPR ratios for TF–target gene (TF–TG) inference using REGA-perturb, PCC, GRNBoost2, SCENIC and CellOracle in K562 GWPS (**f**), K562 essential (**g**) and HepG2 (**h**). ENCODE TF ChIP-seq experiments were used as the benchmark reference (n = 152, 157 and 110 experiments covering 102, 105 and 88 TFs, respectively; panel titles indicate the number of TFs). Statistical significance was assessed using two-sided paired t-tests across experiments (REGA-perturb versus PCC, GRNBoost2, SCENIC and CellOracle: P = 3.5 × 10^−27^, 9.5 × 10^−33^, 6.6 × 10^−38^ and 4.3 × 10^−37^ in **f**; 4.0 × 10^−28^, 5.0 × 10^−37^, 3.8 × 10^−41^ and 8.9 × 10^−36^ in **g**; and 6.5 × 10^−25^, 1.5 × 10^−33^, 1.1 × 10^−35^ and 2.9 × 10^−30^ in **h**; ****P < 0.0001). Box plots within the violins indicate the median and IQR, and the dashed line indicates an AUPR ratio of 1. **i–k**, Precision–recall curves for representative ChIP-seq experiments: E2F4 in K562 GWPS (**i**), MXI1 in K562 essential (**j**) and YY1 in HepG2 (**k**). AUPR ratios for each method are indicated in the legends.

To evaluate the cis-regulatory inferences of REGA-perturb, we used RNA polymerase II ChIA-PET loops from ENCODE^27^ in K562 and HepG2 cells as ground truth. These loops capture chromatin interactions at actively transcribed genes, including enhancer–promoter contacts^32^. We benchmarked the RE–TG weights of REGA-perturb against three alternatives: a PCC baseline, computed as the correlation between RE accessibility and target gene expression in independent paired single-cell multiome data (K562 cells^33^ for the two K562 datasets and primary hepatocytes^34^ for HepG2); a genomic distance decay model; and a random baseline. We did not expect REGA-perturb to surpass the PCC baseline, because PCC is derived from paired single-cell RNA and ATAC profiles, whereas REGA-perturb uses only a reference RE catalog and no quantitative accessibility data. RE–TG pairs were stratified into four distance bins, from 0–50 kb to 200–300 kb relative to the transcription start site (TSS), and performance was measured by the area under the precision–recall curve (AUPR) ratio. In both K562 datasets, REGA-perturb outperformed all baselines in every distance bin, with AUPR ratios of 1.19–1.90 in K562 GWPS and 1.18–1.94 in K562 essential, compared with 0.87–1.21 for the distance baseline and 0.90–0.98 for the PCC baseline (Fig. 2c,d). In HepG2, REGA-perturb outperformed all baselines within 50 kb of the TSS (1.19 versus 1.09 for PCC), whereas, as anticipated, the PCC baseline was higher at longer distances (1.54–2.29 versus 1.39– 1.84 for REGA-perturb; Fig. 2e). When the ground truth was restricted to high-quality loops, defined as those with a paired-end tag count at or above the 75th percentile of each experiment, the AUPR ratios of REGA-perturb increased to 1.25–2.33 in K562 and 1.28–2.00 in HepG2, and the ranking of the methods was unchanged (Supplementary Fig. 1c–e). These results were consistent across complementary evaluation metrics (Supplementary Fig. 1f–k; Supplementary Fig. 2). Thus, in this benchmark, the RE– TG links inferred by REGA-perturb without paired accessibility profiles agreed with chromatin loops better than the baselines at all tested distances in K562 and at promoter-proximal distances in HepG2.

To evaluate trans-regulatory inference, we benchmarked the TF–TG predictions of REGA-perturb against four baselines computed from the same datasets: GRNBoost2^22^, a widely used tree-based method for GRN inference, and a PCC baseline based on the correlation between TF and target gene expression; and two methods that additionally use TF binding motifs, SCENIC^20^ and CellOracle^21^. As ground truth, we used ENCODE TF ChIP-seq data^27^ from the same cell lines, comprising 152, 157 and 110 experiments (102, 105 and 88 TFs) for K562 GWPS, K562 essential and HepG2, respectively. All four baselines performed only marginally better than random, with mean AUPR ratios of 1.16–1.20 for PCC, 1.06–1.12 for GRNBoost2, 1.03–1.06 for SCENIC and 1.11–1.14 for CellOracle. By contrast, REGA-perturb achieved mean AUPR ratios of 1.86 in both K562 datasets and 2.04 in HepG2, which were significantly higher than those of all four baselines (paired two-sided t-test, P < 1 × 10^−24^ for all comparisons; −log10(P) = 24.19– 40.42; Fig. 2f–h) and corresponded to a gain over random prediction more than four times that of the best baseline. For example, for E2F4 in K562 GWPS, MXI1 in K562 essential and YY1 in HepG2, REGA-perturb reached AUPR ratios of 3.37, 3.06 and 3.17, whereas the precision–recall curves of all four baselines remained close to the random expectation (AUPR ratios of 0.84–1.19; Fig. 2i–k). This advantage was consistent across complementary evaluation metrics (Supplementary Fig. 3). Together, these results indicate that REGA-perturb improves the inference of TF–TG relationships over both expression-based and motif-informed methods in all three datasets.

### REGA-perturb distinguishes direct from indirect targets in TF perturbation responses

Knockdown of a TF changes the expression of the genes that it regulates directly as well as genes that respond secondarily^4,5^. We therefore asked whether REGA-perturb can distinguish direct from indirect targets among the genes that respond to a TF perturbation. Following previous studies that integrated TF perturbation responses with ChIP-seq binding to identify direct targets^5,7,35^, we defined putative direct targets of a TF as genes that both responded to knockdown of the TF and showed strong binding by that TF, as quantified by the regulatory potential (RP) score; the remaining differentially expressed genes were considered indirect targets (Methods). Differentially expressed genes were obtained from independent bulk knockdown experiments in KnockTF^36^ for K562 (29 TFs) and from the Perturb-seq screen for HepG2 (15 TFs). For each TF, each method assigned a score to every differentially expressed gene, and we evaluated how well these scores prioritized putative direct targets over indirect targets. REGA-perturb achieved mean AUPR ratios of 1.40 in K562 and 1.73 in HepG2, significantly higher than all four baselines (Fig. 3a,b). In K562, this corresponded to a gain over random prediction about 2.5 times that of the best baseline (1.09 for PCC, 1.11 for GRNBoost2, 1.07 for SCENIC and 1.16 for CellOracle), whereas in HepG2, where the baselines remained close to random (0.96 for PCC, 0.97 for GRNBoost2, 1.01 for SCENIC and 1.08 for CellOracle), the gain was about nine times that of the best baseline.

**Figure 3.**
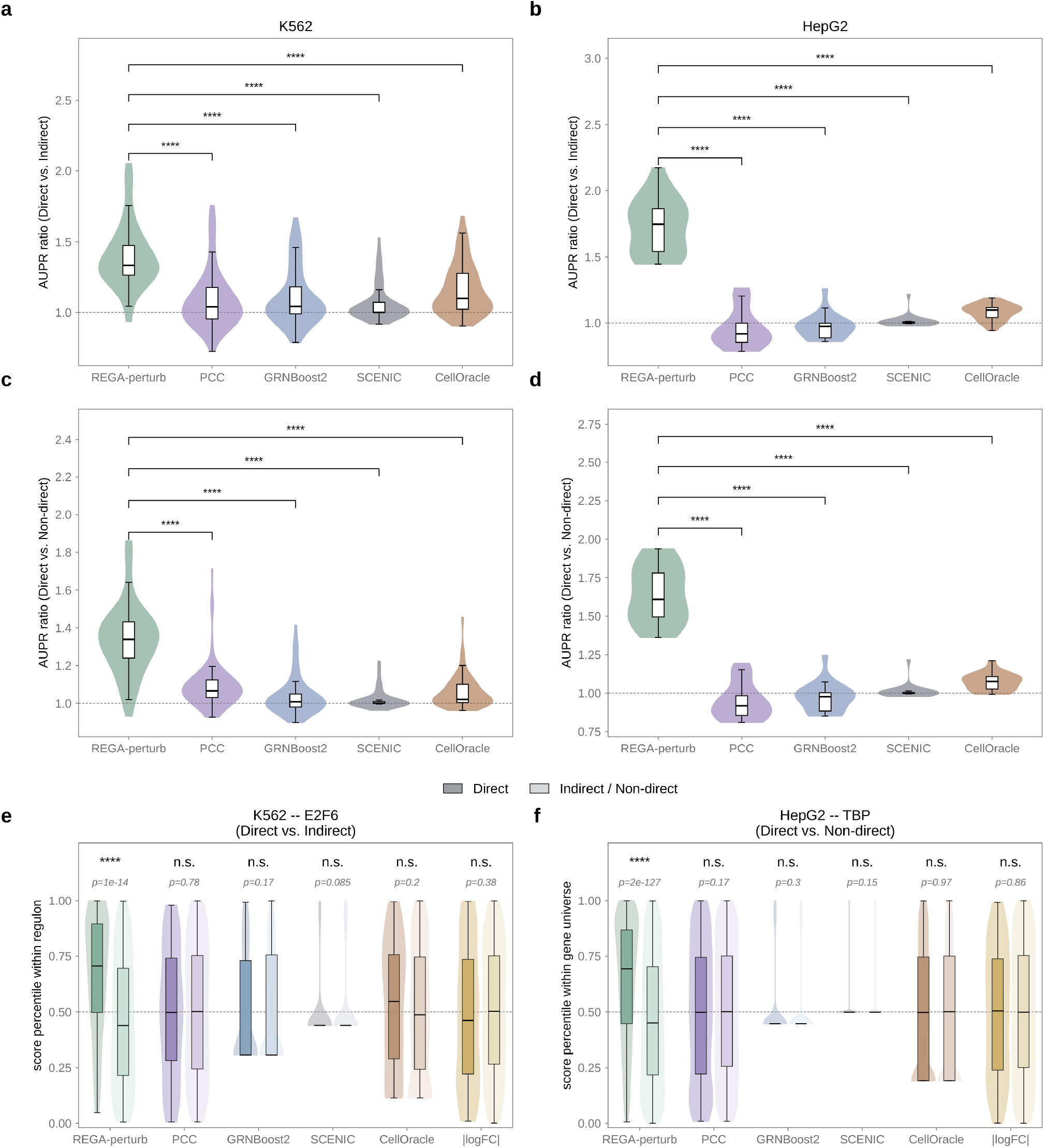
REGA-perturb distinguishes direct from indirect targets in TF perturbation responses. **a–d**, Violin plots comparing AUPR ratios for distinguishing candidate direct targets of transcription factors (TFs) from other genes using REGA-perturb, PCC, GRNBoost2, SCENIC and CellOracle. Candidate direct targets were defined as genes within the top 20% of regulatory potential (RP) scores from TF ChIP-seq experiments. Each violin summarizes TF–ChIP-seq experiment pairs. In K562, pairs were evaluated separately using networks inferred from the essential-gene and genome-wide Perturb-seq screens. **a**,**b**, Discrimination of candidate direct and indirect targets among genes differentially expressed following TF knockdown, using differentially expressed genes (DEGs) from independent bulk knockdown experiments in KnockTF for K562 (**a**; n = 69 pairs from 29 TFs, including 34 from the essential-gene screen and 35 from the genome-wide screen) and pseudobulk Perturb-seq DEGs for HepG2 (**b**; n = 20 pairs from 15 TFs). **c**,**d**, Discrimination of candidate direct and non-direct targets among all genes included in the model for K562 (**c**; n = 67 pairs from 28 TFs) and HepG2 (**d**; n = 20 pairs from 15 TFs). Boxplots within violins indicate the median and interquartile range (IQR), and the dashed line indicates an AUPR ratio of 1. Statistical significance was assessed using two-sided paired Wilcoxon signed-rank tests (REGA-perturb versus PCC, GRNBoost2, SCENIC and CellOracle: P = 7.5 × 10^−9^, 8.7 × 10^−9^, 1.5 × 10^−10^ and 7.3 × 10^−8^ in **a**; 5.5 × 10^−10^, 2.6 × 10^−11^, 7.6 × 10^−12^ and 1.1 × 10^−11^ in **c**; and P = 1.9 × 10^−6^ for all eight comparisons in **b** and **d**, corresponding to the minimum attainable two-sided exact P value for 20 pairs, with REGA-perturb outperforming each baseline in every paired experiment; ****P < 0.0001). **e**,**f**, Violin plots with embedded boxplots showing percentile ranks of gene-level scores for candidate direct (dark) and indirect or non-direct (light) targets, using REGA-perturb, PCC, GRNBoost2, SCENIC, CellOracle and the absolute log fold change following TF knockdown (|logFC|). **e**, E2F6 in the K562 essential-gene screen, comparing direct and indirect genes among those differentially expressed after E2F6 knockdown in the screen (pseudobulk DEGs) (P = 1.4 × 10^−14^ for REGA-perturb, 0.78 for PCC, 0.17 for GRNBoost2, 0.085 for SCENIC, 0.20 for CellOracle and 0.38 for |logFC|). **f**, TBP in HepG2, comparing direct and non-direct genes among all genes included in the model (P = 2.0 × 10^−127^, 0.17, 0.30, 0.15, 0.97 and 0.86, respectively). The dashed line indicates the 50th percentile; P values above each pair are from two-sided Mann–Whitney U tests (n.s., P ≥ 0.05; ****P < 0.0001).

To test whether this result depended on restricting the analysis to differentially expressed genes, we repeated the analysis using all genes in the model as the background, comparing the genes with the highest RP scores with all remaining genes. REGA-perturb again achieved significantly higher AUPR ratios than all four baselines, with mean AUPR ratios of 1.34 in K562 and 1.64 in HepG2, compared with 0.95–1.09 for the baselines (Fig. 3c,d). The advantage of REGA-perturb was robust to both the definition of direct targets and the evaluation metric: the results were consistent when direct targets were additionally required to be supported by a ChIA-PET loop and when performance was assessed by AUROC (Supplementary Fig. 4a–l). The same pattern was also observed when the K562 analysis was repeated using differentially expressed genes from the Perturb-seq screens, with the two screens analyzed separately (Supplementary Fig. 5).

This advantage was also evident at the level of individual TFs. For example, among genes differentially expressed after E2F6 knockdown in K562 essential, TF–TG scores from REGA-perturb were significantly higher for putative direct targets than for indirect targets (P = 1.4 × 10^−14^), whereas scores from PCC, GRNBoost2, SCENIC and CellOracle did not differ between the two groups (P = 0.78, 0.17, 0.085 and 0.20, respectively). The magnitude of the expression response itself, measured as the absolute log fold change (|logFC|) after knockdown, likewise did not distinguish direct from indirect targets (P = 0.38; Fig. 3e). The same pattern was observed for TBP in HepG2 when all genes in the model were used as the background, with REGA-perturb significantly separating the two groups (P = 2.0 × 10^−127^), whereas the other scores did not (P ≥ 0.15; Fig. 3f). Similar results were observed for two additional K562 TFs, BHLHE40 and STAT1 (Supplementary Fig. 4m,n). Together, these results indicate that REGA-perturb more effectively prioritizes ChIP-supported direct targets over indirect targets in TF perturbation responses than both expression-based and motif-informed methods.

### REGA-perturb predicts regulons responsive to unseen TF perturbations

Most perturbation screens cover only a subset of TFs in a given cell type, and a regulatory network is more useful if it can anticipate the effects of perturbations that have not been performed. We therefore asked whether REGA-perturb can predict regulons responsive to unseen TF perturbations. We retrained REGA-perturb using only data from non-TF perturbations, and applied the baseline methods to the same data. SCENIC was not included in this comparison because it failed to yield a regulon for a large fraction of the unseen TFs. For each unseen TF, we defined its predicted regulon as the genes with the highest TF– TG scores and its regulon activity as the mean expression of these genes for each cell (Methods).

For each unseen TF, the activity of the regulons predicted by REGA-perturb distinguished cells in which that TF was perturbed from control cells. The average AUPR ratios were 2.07, 2.08 and 2.04 for the 35, 262 and 41 unseen TFs in K562 essential, K562 GWPS and HepG2, respectively, significantly higher than all three baselines (1.38–1.62; paired two-sided t-test, P < 2 × 10^−3^ for all comparisons; −log10(P) = 2.92– 26.56; Fig. 4a–c). The advantage of REGA-perturb was consistent when performance was assessed by AUROC (Supplementary Fig. 6a–c). For FOXM1 in K562 GWPS, for example, the activity of the regulon predicted by REGA-perturb differed significantly between FOXM1 knockdown and control (two-sided Mann–Whitney U test, P = 7.4 × 10^−4^; Fig. 4d), whereas the activities of the regulons predicted by PCC, GRNBoost2 and CellOracle did not differ significantly (P = 0.66, 0.67 and 0.41, respectively; Fig. 4e–g). Similar results were observed for BDP1 in K562 essential, STAT1 in K562 GWPS and ATF5 in HepG2 (Supplementary Fig. 6d–f). The weaker performance of PCC, GRNBoost2 and CellOracle may reflect their reliance on covariation between the expression of a TF and its target genes, which depends on variation in the expression of the TF and may therefore be less informative when the perturbation of the TF is withheld. Together, these results indicate that REGA-perturb predicts regulons responsive to unseen TF perturbations more accurately than both expression-based and motif-informed methods, suggesting that it can extend regulatory inference to TFs whose perturbations have not been performed.

**Figure 4.**
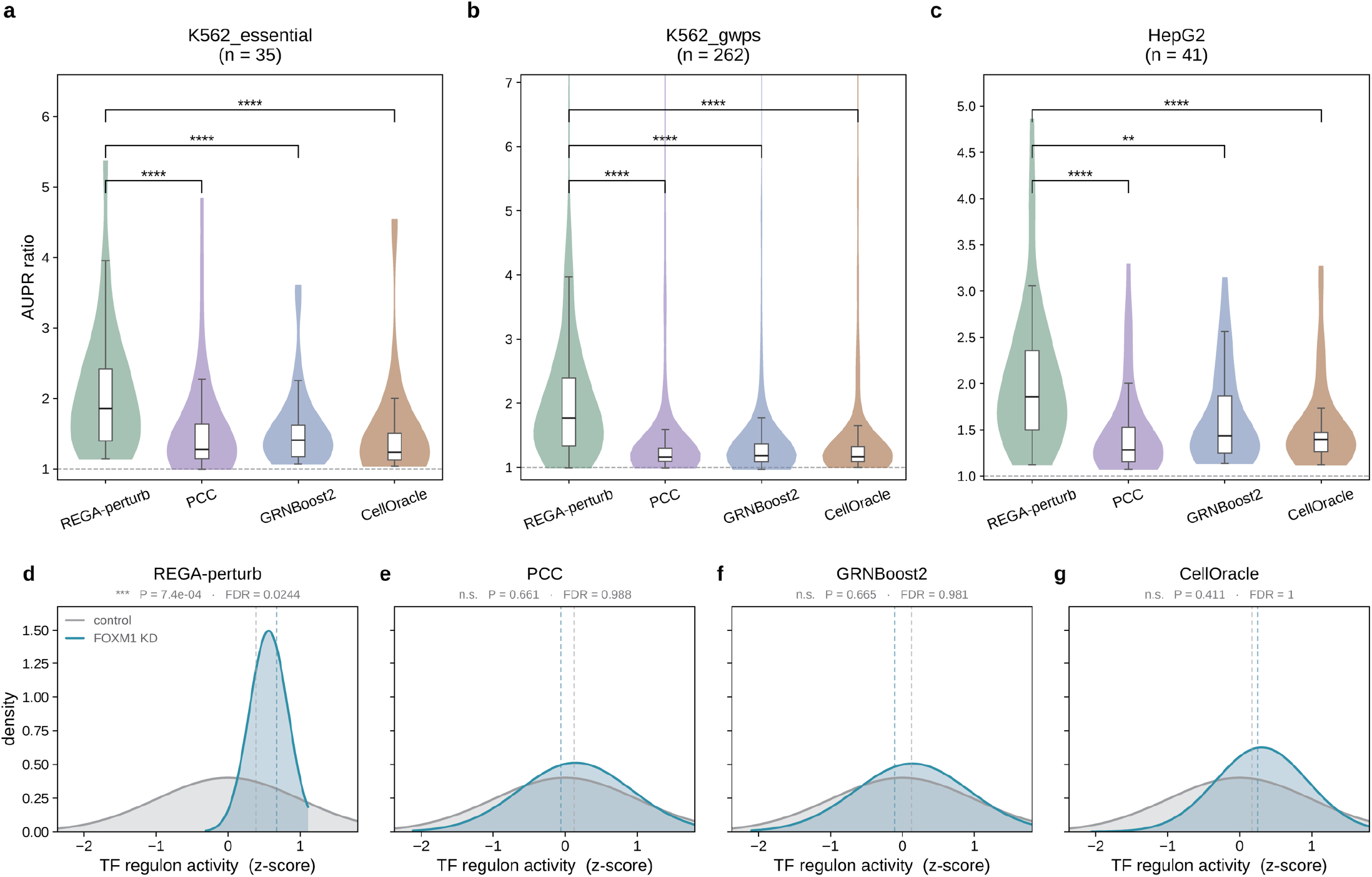
REGA-perturb predicts regulons responsive to unseen TF perturbations. **a–c**, Violin plots comparing area under the precision–recall curve (AUPR) ratios for distinguishing knockdown from control cells based on regulon activity for transcription factors (TFs) whose perturbations were excluded from model training (unseen TFs). Results are shown for the K562 essential-gene screen (K562 essential; **a**, n = 35 unseen TFs), K562 genome-wide screen (K562 GWPS; **b**, n = 262) and HepG2 (**c**, n = 41). For each unseen TF, regulons were defined as the top 200 genes ranked by REGA-perturb, PCC, GRNBoost2 or CellOracle, and regulon activity was calculated as the mean expression of these genes. Paired comparisons were performed using the unseen TFs shared by REGA-perturb and each baseline (n = 34, 249 and 37 for CellOracle in **a, b** and **c**, respectively). Boxplots within violins indicate the median and interquartile range (IQR), and the dashed line indicates an AUPR ratio of 1. The y axis is truncated at the pooled 99th percentile. To account for potential increases or decreases in regulon activity following TF knockdown, each AUPR ratio was calculated as the larger of the values obtained by ranking regulon activity in ascending or descending order. Statistical significance was assessed using two-sided paired t-tests across unseen TFs (REGA-perturb versus PCC, GRNBoost2 and CellOracle: P = 5.9 × 10^−6^, 1.1 × 10^−5^ and 6.6 × 10^−7^ in **a**; 2.7 × 10^−27^, 2.4 × 10^−24^ and 4.0 × 10^−23^ in **b**; and 1.7 × 10^−5^, 1.2 × 10^−3^ and 6.7 × 10^−5^ in **c**; **P < 0.01, ****P < 0.0001). **d–g**, Representative example of an unseen TF, FOXM1, in K562 GWPS. Density plots showing regulon activity, expressed as z-scores relative to controls, in knockdown and control cells for regulons predicted by REGA-perturb (**d**), PCC (**e**), GRNBoost2 (**f**) and CellOracle (**g**). Curves represent Gaussian fits, and dashed lines indicate group medians. P values were calculated using two-sided Mann–Whitney U tests, with false discovery rates (FDRs) estimated using the Benjamini– Hochberg procedure across all unseen TFs evaluated by each method (P = 7.4 × 10^−4^, FDR = 0.024 in **d**; P = 0.66, FDR = 0.99 in **e**; P = 0.67, FDR = 0.98 in **f**; and P = 0.41, FDR = 1.00 in **g**; n.s., P ≥ 0.05; ***P < 0.001).

### REGA-perturb captures perturbation-induced changes in TF activity

Beyond identifying the downstream targets of individual TFs, understanding cellular responses to perturbations requires characterizing their effects on other regulators. REGA-perturb links each perturbation target directly to TFs through the regulatory path of the model, connecting perturbation targets to regulatory programs and subsequently to TF motifs (Fig. 1a). By integrating these relationships, REGA-perturb estimates TF activity for each perturbation–TF pair by combining the loadings of the TF’s motifs on the latent regulatory programs with the program activities of the perturbation, and quantifies perturbation-induced changes relative to non-targeting control cells (Methods). We investigated whether these inferred changes capture the regulatory consequences of perturbations.

Perturbations with strong transcriptomic effects changed the inferred activity of many TFs, as illustrated by the most strongly perturbed TFs in each dataset (Fig. 5a; Supplementary Fig. 7a,b). For example, GATA1 knockdown, which exhibited the greatest perturbation strength in K562 essential, also produced the largest changes in inferred TF activity (Fig. 5a). Knockdowns of functionally related regulators clustered together, including MAX and its interaction partner MYC^37^, as well as CTCF and the cohesin subunits SMC3 and RAD21^38^, suggesting that they affected TF activity in similar ways (Fig. 5a). Whereas knockdown reduces the expression of the targeted TF, REGA-perturb infers TF activity at the regulatory level. Stronger knockdowns led to larger decreases in the inferred activity of the knocked-down TF itself, and all TFs whose own activity changed substantially, such as GATA1, SMC3, CDC5L and MAX, showed decreases (Fig. 5b). Similar but weaker trends were observed in K562 GWPS and HepG2 (Supplementary Fig. 7c,d).

**Figure 5.**
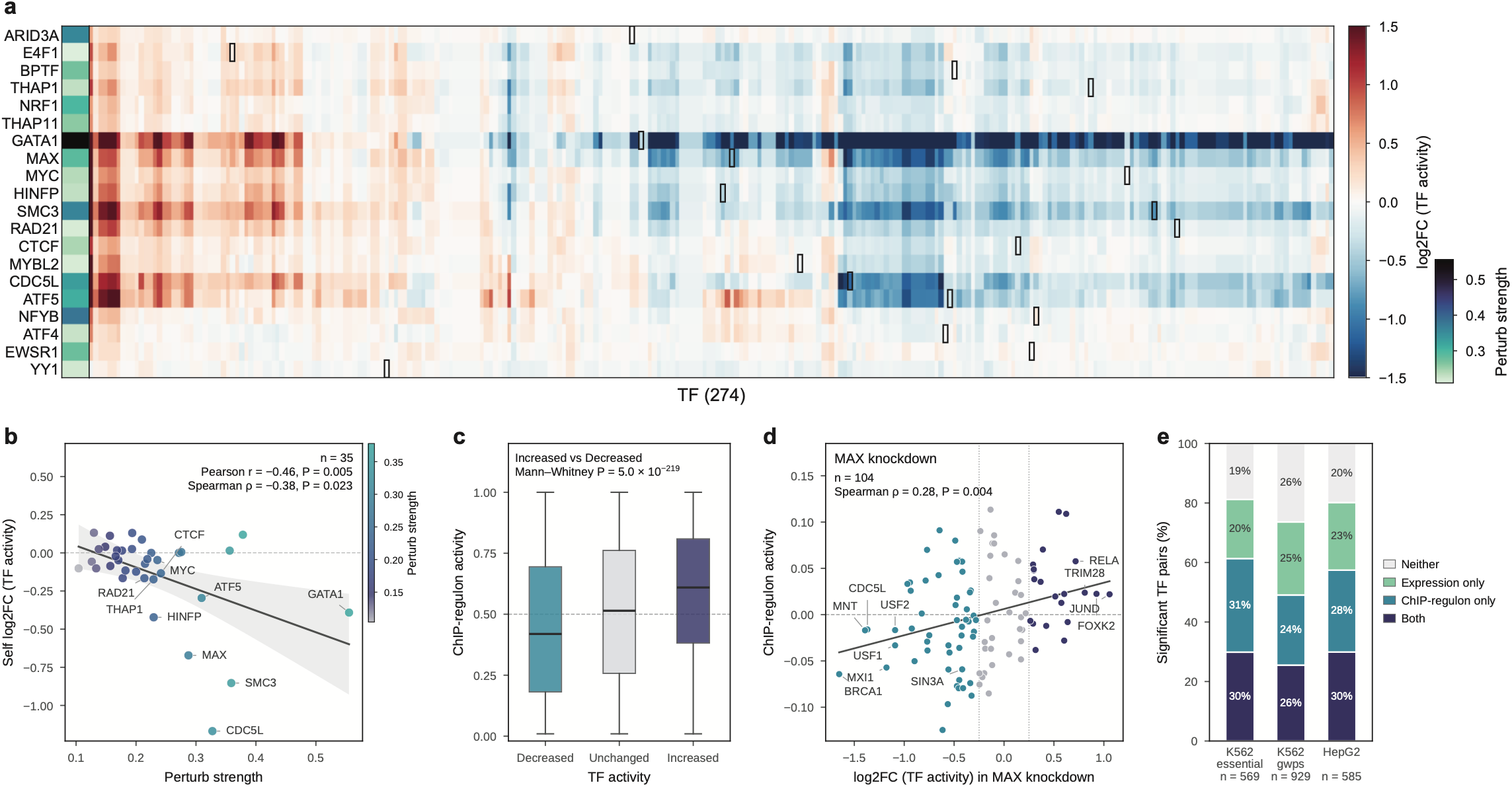
REGA-perturb captures perturbation-induced changes in TF activity. **a**, Heatmap showing log_2_ fold changes (log_2_FCs) in inferred transcription factor (TF) activity relative to non-targeting controls for 274 modeled TFs (columns) following knockdown of the 20 TFs with the strongest perturbation effects (rows) in the K562 essential-gene screen (K562 essential). Perturbation strength (left color bar) was defined as the root mean square of gene-level log fold changes (logFCs). Rows and columns were hierarchically clustered using average linkage and correlation distance. The color scale is limited to ±1.5, and black boxes mark the perturbed TF. **b**, Association between perturbation strength and changes in the perturbed TF’s own inferred activity for 35 TFs in K562 essential. Points are colored by perturbation strength. The solid line shows the least-squares fit with its 95% confidence interval, and the dashed line indicates no activity change (two-sided correlation tests; Pearson r = −0.46, P = 0.005; Spearman ρ = −0.38, P = 0.023). The ten TFs with the highest combined ranks for perturbation strength and activity decrease are labeled. **c**, Boxplots comparing ChIP-regulon activity among TFs predicted to have decreased (log_2_FC ≤ −0.25; n = 12,129 knockdown–TF pairs), unchanged (|log_2_FC| < 0.25; n = 33,334) or increased (log_2_FC ≥ 0.25; n = 7,030) activity across the 500 strongest perturbations in K562 essential, excluding the perturbed TF itself. Each ChIP regulon comprised the top 200 genes ranked by regulatory potential scores from ENCODE TF ChIP-seq experiments. ChIP-regulon activity was defined as the mean logFC of regulon genes, averaged across ChIP-seq experiments for each TF and expressed as a percentile among TFs within each knockdown. Boxes indicate the median and interquartile range (IQR), whiskers extend to 1.5 × IQR, outliers are omitted, and the dashed line indicates 0.5. Median activities were 0.42, 0.51 and 0.61, respectively (two-sided Mann–Whitney U test comparing increased and decreased groups, P = 5.0 × 10^−219^). **d**, Representative example of MAX knockdown in K562 essential. Scatter plot comparing predicted TF activity changes (log_2_FC) with ChIP-regulon activity for 104 TFs with available ChIP-seq data. ChIP-regulon activity was calculated as the mean logFC of each TF’s ChIP regulon, centered by subtracting the mean across TFs within the MAX knockdown. Colors correspond to the predicted activity categories in **c**. Dotted lines indicate log_2_FC thresholds of ±0.25, and the solid line represents the least-squares regression fit. A positive association was observed between predicted TF activity changes and ChIP-regulon activity (two-sided Spearman correlation, ρ = 0.28, P = 0.004). **e**, Stacked bar plots showing the agreement between predicted TF activity changes and two independent measures of transcriptional responses: changes in ChIP-regulon activity and changes in the TF’s own expression. Analyses included knockdown–TF pairs with predicted activity changes (|log_2_FC| ≥ 0.25) among the 20 strongest perturbations in each dataset. Pairs were classified according to whether the predicted direction agreed with both measures, only one measure or neither. Results are shown for K562 essential (n = 569), K562 GWPS (n = 929) and HepG2 (n = 585).

To evaluate these inferred TF activity changes, we compared them with the expression changes of the target genes of each TF defined by independent ChIP-seq data (ChIP-regulon activity; Methods). TFs predicted to increase in activity had significantly higher ChIP-regulon activity than those predicted to decrease (two-sided Mann–Whitney U test, P = 5.0 × 10^−219^ in K562 essential; Fig. 5c). For example, after MAX knockdown, the predicted changes in TF activity correlated with ChIP-regulon activity (Spearman correlation, ρ = 0.28, P = 0.004), and the inferred activities of the MAX-dependent repressor MXI1^39^ and its co-repressor SIN3A^40,41^ decreased together with the expression of their ChIP-seq target genes (Fig. 5d). Similar results were observed in K562 GWPS and HepG2 (Supplementary Fig. 7e,f). To further assess agreement with expression data not used for training, we repeated the analysis using independent knockdown experiments in K562 from KnockTF and observed the same trend (Supplementary Fig. 7g,h). Consistent with activity being inferred at the regulatory level, the predicted direction of change agreed with the change in the TF’s own expression in only about half of knockdown–TF pairs, and it agreed with ChIP-regulon activity, TF expression or both in 81%, 74% and 80% of pairs in K562 essential, K562 GWPS and HepG2, respectively (Fig. 5e). Together, these results indicate that REGA-perturb infers perturbation-induced changes in TF activity that are consistent with changes in the expression of ChIP-seq target genes but are not simply reflected by changes in the expression of the TFs themselves.

### REGA-perturb resolves perturbation-specific regulatory rewiring

Genes directly bound by a perturbed TF might be expected to show the largest expression responses. Contrary to this expectation, expression changes following TF perturbation are often only weakly associated with TF binding measured by ChIP-seq^5,7^. We observed this negative association in KnockTF knockdown experiments: across 118 experiments targeting 96 TFs in K562, absolute gene expression changes showed negative correlation with the ChIP-seq regulatory potential (RP) scores of the perturbed TFs (median Spearman’s ρ = −0.02; two-sided Wilcoxon signed-rank test against shuffled RP scores, P = 8.5 × 10^−3^; Supplementary Fig. 8d). For example, following ZBTB33 knockdown, absolute gene expression changes were negatively correlated with ZBTB33 RP scores (Spearman’s ρ = −0.07, P = 5.7 × 10^−9^; Fig. 6a). Similarly negative associations were observed across the three Perturb-seq screens, including 300 perturbed TFs in K562 GWPS (median ρ = −0.03; Fig. 6b; Supplementary Fig. 8a–c). Thus, the genes showing the largest transcriptional responses were not necessarily those most strongly bound by the perturbed TF.

**Figure 6.**
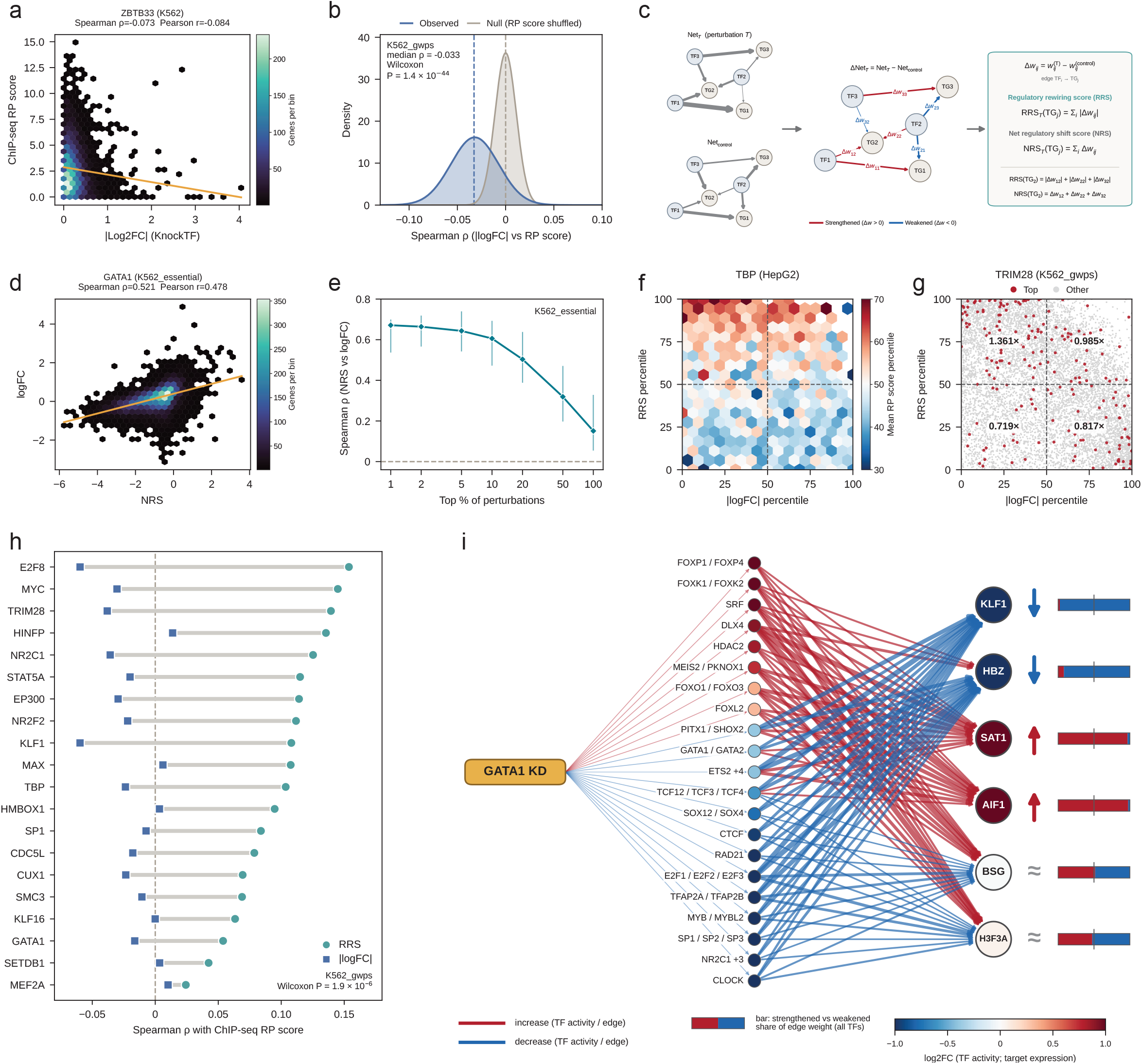
REGA-perturb resolves perturbation-specific regulatory rewiring. **a**, Hexbin plot showing the relationship between the ChIP-seq regulatory potential (RP) scores of ZBTB33 and absolute gene expression changes (|log_2_FC|) following ZBTB33 knockdown in K562, using an independent bulk knockdown experiment from KnockTF (n = 6,332 genes). Color indicates gene density, and the line represents the least-squares fit (Spearman ρ = −0.073, two-sided P = 5.7 × 10^−9^). **b**, Distributions of Spearman correlations between absolute expression changes (|logFC|) following knockdown and the RP scores of the perturbed transcription factors (TFs) across genes for 300 TF perturbations with ChIP-seq data in the K562 genome-wide screen (K562 GWPS), compared with correlations obtained by shuffling RP scores across genes. Curves represent normal fits, and dashed lines indicate medians (two-sided paired Wilcoxon signed-rank test, P = 1.4 × 10^−44^). **c**, Schematic of perturbation-specific regulatory rewiring. For each perturbation, the differential regulatory network was defined by subtracting the control network from the perturbation-specific network. Changes in TF–target gene (TF–TG) edge weights (Δw) indicate increased (red) or decreased (blue) inferred regulatory influence. For each target gene, the regulatory rewiring score (RRS) was calculated as the sum of |Δw|, whereas the net regulatory shift score (NRS) was calculated as the sum of Δw across incoming TF edges. **d**, Hexbin plot showing the relationship between NRS and gene expression changes (logFC) following GATA1 knockdown in the K562 essential-gene screen (K562 essential; n = 8,535 genes; Spearman ρ = 0.521, two-sided P < 1 × 10^−300^). **e**, Spearman correlations between NRS and logFC across genes for increasing proportions (top 1–100%; n = 20–2,046) of K562 essential perturbations ranked by the magnitude of their inferred regulatory changes, measured as the Euclidean distance between perturbation-specific and control regulatory program activity profiles. Points indicate medians, and vertical lines indicate interquartile ranges. **f**, Hexbin plot showing the percentile ranks of absolute expression changes (|logFC|; x axis) and RRS (y axis) following TBP knockdown in HepG2 (n = 9,607 genes). Color indicates the mean percentile rank of TBP ChIP-seq RP scores within each bin, and dashed lines indicate medians. **g**, Scatter plot showing the corresponding relationship following TRIM28 knockdown in K562 GWPS (n = 8,221 genes). The 200 genes with the highest TRIM28 RP scores are highlighted in red, with fold enrichment indicated for each quadrant (one-sided binomial test, n = 200 genes; P = 5.1 × 10^−4^ for the high-RRS, low-|logFC| quadrant). **h**, Dumbbell plot comparing Spearman correlations of RRS (circles) and |logFC| (squares) with the RP scores of the perturbed TFs across genes for 20 TF perturbations in K562 GWPS (two-sided paired Wilcoxon signed-rank test, n = 20 TFs, P = 1.9 × 10^−6^). These TFs had annotated motifs, showed decreased expression following their own knockdown and ranked highest in inferred regulatory changes, as defined in **e. i**, Differential regulatory network for six target genes following GATA1 knockdown in K562 essential. TF nodes represent TFs with predicted activity changes of |log_2_FC| ≥ 0.25 and are colored by their inferred activity changes. TFs sharing identical activity changes and edge weights owing to shared motifs were merged into single nodes, labeled with up to three TF names or the first name followed by the number of additional TFs (for example, ETS2 +4). Target genes are colored by their expression log_2_FCs, with arrows indicating decreased, increased or nearly unchanged expression. Edges indicate increases (red) or decreases (blue) in inferred regulatory influence, with widths proportional to |Δw| relative to the largest edge for each gene. For visualization, only the eight TF nodes with the largest |Δw| were displayed for each gene, except for genes with nearly unchanged expression, for which the eight largest positive and eight largest-magnitude negative Δw values were selected separately. Edges were retained if |Δw| reached at least 25% of the maximum |Δw| for each gene, with the threshold reduced to 15% for genes with nearly unchanged expression to preserve both strengthened and weakened interactions. These thresholds were applied only to the visualization; the bars indicate the proportions of strengthened and weakened edge weights calculated across all modeled TFs for each target gene. Thin lines connecting the perturbation to TF nodes indicate the direction of predicted TF activity changes.

This apparent disconnect may arise because gene expression reports only the net outcome of multiple regulatory inputs. Following perturbation, some inputs to a gene may strengthen while others weaken, causing their effects to cancel and leaving expression nearly unchanged despite extensive reorganization of the gene’s regulatory architecture. Although regulatory compensation is a fundamental feature of biological systems^18,42^, it has been difficult to quantify directly from perturbation-induced expression changes. REGA-perturb addresses this limitation by inferring how each TF–gene interaction changes following perturbation. For each perturbation, we constructed a differential regulatory network in which each TF–gene edge represents its change in weight relative to non-targeting control cells (Fig. 6c). We then defined two complementary gene-level scores: the net regulatory shift score (NRS), the signed sum of changes in incoming edges, captures the net regulatory effect after opposing inputs cancel; the regulatory rewiring score (RRS), the sum of their absolute changes, captures the total magnitude of regulatory change before cancellation (Fig. 6c; Methods).

The NRS showed strong correlations with observed expression changes, with median Spearman’s ρ ranging from 0.51 to 0.79 among the top 1% of perturbations across the three datasets (Fig. 6e; Supplementary Fig. 8e,f). For example, following GATA1 knockdown in K562 essential, gene-level NRS values correlated with the direction and magnitude of expression changes (Spearman’s ρ = 0.52, P < 10^−300^; Fig. 6d). Whereas the NRS summarizes the net regulatory shift, the RRS captures the total magnitude of regulatory changes. We therefore examined whether the RRS was more closely associated with binding of the perturbed TF than absolute expression changes. Across nearly all perturbed TFs with available ChIP-seq data, the RRS showed stronger correlations with the RP scores of the perturbed TFs than did absolute expression changes (87 of 88, 297 of 300 and 63 of 63 TFs in K562 essential, K562 GWPS and HepG2, respectively; median ρ = 0.09–0.12 versus −0.03 to −0.02; two-sided paired Wilcoxon signed-rank test, P < 10^−11^ for all comparisons; −log10(P) = 11.29–50.15; Supplementary Fig. 9c–h). For example, among 20 selected TFs in K562 GWPS, the RRS was more strongly correlated with RP scores than absolute expression changes for every TF (median ρ = 0.11 versus −0.02; two-sided paired Wilcoxon signed-rank test, P = 1.9 × 10^−6^; Fig. 6h), with similar results in K562 essential and HepG2 (Supplementary Fig. 9a,b). This advantage persisted when expression changes were obtained from independent KnockTF knockdown experiments (Supplementary Fig. 9i–l).

Genes bound by the perturbed TF were enriched among genes with high RRS but not among genes with large expression changes. Across all perturbed TFs, genes with the highest RP scores were enriched in the upper half of the RRS distribution (median 1.18-, 1.19- and 1.23-fold in K562 essential, K562 GWPS and HepG2, respectively; Supplementary Fig. 8i–k), whereas absolute expression changes showed little correlation with RP scores (Fig. 6b). For example, TBP-bound genes in HepG2 were concentrated in the upper half of the RRS distribution regardless of their expression changes (Fig. 6f). Similarly, TRIM28-bound genes in K562 GWPS were enriched among genes with high RRS but small expression changes (1.36-fold; one-sided binomial test, P = 5.1 × 10^−4^; Fig. 6g). Enrichment of TRIM28-bound genes among high-RRS genes was also observed using expression changes from an independent TRIM28 knockdown experiment (Supplementary Fig. 8g,h).

The differential regulatory networks further revealed distinct patterns of regulatory rewiring across genes following the same perturbation (Fig. 6i). Following GATA1 knockdown in K562 essential, nearly all incoming edges of the erythroid genes KLF1 and HBZ, both known GATA1 targets^43,44^, were weakened, consistent with their reduced expression. These included the GATA1/GATA2-to-KLF1 edge, accompanied by decreased inferred activity of GATA1/GATA2. In contrast, the incoming edges of SAT1 and AIF1 were strengthened, consistent with their increased expression. For BSG and H3F3A, strengthened and weakened edges largely offset one another, and expression remained nearly unchanged despite substantial regulatory rewiring. Similar patterns were observed following CEBPG knockdown in HepG2 and MYB knockdown in K562 GWPS (Supplementary Fig. 9m,n). For example, MYB knockdown increased the expression of the ζ-globin gene HBZ, which is repressed by MYB^45^, accompanied by strengthening of nearly all its incoming edges. Together, these results show that expression change and regulatory change represent distinct layers of the perturbation response. The NRS captures the net regulatory shift reflected in gene expression, whereas the RRS captures the total reorganization of regulatory inputs, including opposing changes that can cancel at the expression level. Consequently, genes with little expression change can nevertheless undergo extensive inferred regulatory rewiring. By recovering the binding footprint of the perturbed TF more consistently than differential expression, REGA-perturb provides a framework for detecting—and potentially quantifying—regulatory compensation that is hidden from conventional perturbation analyses.

## Discussions and conclusions

Perturb-seq responses reflect regulatory mechanism, but because each gene’s expression change mixes the contributions of many regulatory paths, the mechanism cannot be resolved from the response alone, however large the model. REGA-perturb addresses this by supplying information that expression data cannot provide. Reference epigenomes define a scaffold of possible TF–RE–gene paths, and responses to thousands of perturbations determine how strongly each path is used and how its use changes from one perturbation to another. Because the inferred paths make explicit mechanistic claims, we evaluated them against independent measurements of regulation rather than by prediction accuracy alone. Across three CRISPRi screens, REGA-perturb inferred TF–target relationships that agreed with TF binding far better than existing methods, prioritized direct over indirect targets, generalized to TFs withheld from training, and resolved perturbation-specific changes in TF activity and in individual regulatory inputs.

These gains likely reflect how the scaffold and the perturbation data compensate for each other’s weaknesses. Motifs and RE–gene proximity define far more candidate interactions than are functional in any given cell type. Expression covariation, on which SCENIC, CellOracle and GRNBoost2 rely, confounds direct regulation with co-regulation and indirect effects. Perturbations resolve part of this ambiguity because they are interventions, and pooling thousands of them through shared programs selects, among the candidate paths, those that consistently explain the observed responses. The two components play distinct roles. The scaffold supplies information external to expression, restricting inference to paths that can exist. The shared programs allow thousands of perturbations to inform the same set of path weights. SCENIC and CellOracle also use motifs, yet remained close to random in our benchmarks, suggesting that the improvement comes not from the motif prior alone but from combining it with perturbation responses. The same pooling may explain why REGA-perturb generalized to TFs withheld from training: a TF’s regulatory paths can be informed by perturbations of other genes that alter its activity. Covariation-based methods, by contrast, depend on variation in the TF’s own expression, which is largely absent when its perturbation is withheld. Notably, these gains were achieved with a linear model, consistent with the view that resolving mechanism depends more on supplying the right information than on model capacity^13^. Methods that integrate single-cell chromatin accessibility, such as LINGER and SCENIC+^25,26^, offer another way to add information beyond expression, but they require accessibility profiles that large-scale Perturb-seq screens lack. Because REGA-perturb requires only RNA, it can be applied directly to existing screens.

Most genes targeted in a genome-wide screen are not TFs, yet many of their effects are likely mediated by TFs. By linking perturbations to TFs through shared programs, REGA-perturb estimates how each perturbation alters the activity of each TF. These estimates agreed with expression changes in independently defined ChIP-seq regulons but matched changes in the TF’s own transcript in only about half of cases, consistent with regulation of TF activity through cofactors, complexes and post-transcriptional mechanisms^14–16^. For example, MAX knockdown reduced the inferred activity of MXI1 and its co-repressor SIN3A, consistent with the requirement of MAX for MXI1-mediated repression^39–41^. Such estimates can help identify the regulators through which a perturbation acts, and can prioritize candidate TFs for follow-up when the perturbed gene is not itself a TF.

A central observation of this study is that expression change and regulatory change are distinct layers of the perturbation response. The same mixing of regulatory inputs that prevents mechanism from being read directly from expression can also conceal regulatory change altogether. Genes bound by a perturbed TF often change little in expression, a pattern usually attributed to nonfunctional binding, context dependence or redundancy^5–7,46^. Our results suggest an additional, not mutually exclusive, explanation: some of these genes undergo substantial changes in regulatory inputs that offset one another. The total magnitude of inferred regulatory change tracked binding of the perturbed TF more closely than absolute expression change did, for 447 of 451 perturbed TFs across three datasets, although the correlations were modest in absolute terms. Such offsetting changes are consistent with regulatory compensation through feedback and redundancy^17,42^. Because successful buffering preserves expression, differential expression analysis classifies these genes as unaffected. Yet a gene whose expression is maintained by rebalanced inputs is not necessarily regulated as before, and may become more dependent on the compensating regulators. Combinatorial perturbation of a primary target together with its predicted compensating regulators^47^ provides a direct test of this prediction and may reveal genetic interactions invisible to single-gene screens.

Several limitations should be noted. First, REGA-perturb relies on a reference RE catalog from unperturbed cells, together with TF motifs and proximity-based RE–gene candidates. It cannot capture regulation through REs absent from the catalog, TFs without known motifs or interactions beyond the proximity window. It also does not model perturbation-induced changes in chromatin accessibility, so the inferred rewiring reflects changes in the use of existing paths rather than the creation of new ones. Second, because candidate TF–RE links are restricted to motif occurrences, motif-based inference and binding-based evaluation are not fully independent. The near-random performance of motif-informed baselines argues that the prior alone does not account for our results, but this dependence should be considered when interpreting absolute performance. Third, the model is linear and represents each perturbation by a single profile of program activities. It therefore does not capture nonlinear interactions between regulators, or heterogeneity among cells receiving the same perturbation. Fourth, inferred changes in TF activity and regulatory inputs are model-based estimates derived from transcriptional data. They do not by themselves establish changes in physical binding or causal mediation through specific REs, which will require targeted RE perturbations, binding measurements after perturbation and time-resolved designs. Finally, we evaluated REGA-perturb in CRISPRi screens of two extensively annotated cell lines. Its performance in primary cells and tissues, where reference epigenomes are sparser, and with other perturbation modalities remains to be established.

Perturb-seq screens are expanding to more cell types, perturbation modalities and molecular readouts^47–50^, and reference epigenomic atlases are expanding in parallel^27,51,52^. As these resources grow, evaluating perturbation models against independent measurements of regulatory mechanism, rather than prediction accuracy alone, will be important for determining whether they capture how perturbations act. REGA-perturb provides a way to bring perturbation screens and reference epigenomes together, turning screens from catalogs of transcriptional responses into maps of the regulatory paths through which perturbations act, and of how those paths are reorganized from one perturbation to another.

## Methods

### Data collection and preprocessing

Perturb-seq data were obtained from three published CRISPR interference (CRISPRi)^53^ screens: genome-wide (K562 GWPS) and essential-gene (K562 essential) screens in K562 cells from Replogle et al.^3^, and a screen in HepG2 cells from Nadig et al.^29^. We used the single-cell unique molecular identifier (UMI) count matrices and corresponding perturbation target annotations provided by the original studies, with cells carrying non-targeting guides serving as controls. Cells with 1,000–50,000 UMIs were retained, and mitochondrial genes and genes detected in fewer than 30 cells were excluded. Following quality control, K562 GWPS contained 1,971,958 cells across 9,866 perturbation targets, including 74,698 non-targeting control cells, with 8,231 genes retained; K562 essential contained 301,774 cells across 2,057 perturbation targets, including 10,476 control cells, with 8,550 genes retained; and HepG2 contained 141,560 cells across 2,393 perturbation targets, including 4,825 control cells, with 9,609 genes retained. Gene expression counts were library-size normalized and log-transformed, and the resulting gene-by-cell expression matrices were used as model inputs.

Reference regulatory element (RE) catalogs were constructed from bulk ATAC-seq data for the corresponding cell lines (K562 and HepG2; GRCh38) obtained from ENCODE^27^. For each cell line, ATAC-seq peaks from multiple experiments were pooled, and overlapping peaks were merged. TF ChIP-seq data were also obtained from ENCODE, comprising 423 experiments covering 304 TFs in K562 and 160 experiments covering 135 TFs in HepG2. For each ChIP-seq experiment, gene-level regulatory potential (RP) scores were calculated using the BETA model^35^. RNA polymerase II (POLR2A) ChIA-PET loops and H3K27ac and H3K4me3 ChIP-seq peaks for both cell lines were also obtained from ENCODE. Base-wise evolutionary conservation scores (phastCons^54^, 100-way vertebrate alignment; hg38) were downloaded from the UCSC Genome Browser^55^. Bulk TF knockdown and knockout data for K562 cells were obtained from KnockTF 2.0^36^. Paired single-nucleus RNA-seq and ATAC-seq profiles of K562 cells generated using SNARE-seq^33^, together with single-nucleus multiome data of hepatocytes from human liver tissue^34^, were used to calculate RE–gene correlation baselines.

### REGA-perturb model

REGA-perturb models the effect of a genetic perturbation on single-cell gene expression by explicitly propagating perturbation signals through a hierarchical regulatory cascade. Each perturbation target is first represented by its activity over a small set of latent regulatory programs, which capture shared modes of perturbation response. These programs are linked to transcription factor (TF) motifs, whose activities are propagated to regulatory elements (REs), and then from REs to their downstream target genes. In this way, the model connects each perturbation target to the observed expression profile of a cell through the chain perturbation target → regulatory programs → TF motifs → REs → target genes (Fig. 1a).

A central component of REGA-perturb is the reference regulatory path, which defines the downstream segment of this cascade from TF motifs to REs and from REs to genes. This path is derived from the reference RE catalog constructed for the matched cell type and encodes prior regulatory knowledge, including motif occurrence within REs and candidate RE–target gene links. By constraining how perturbation-induced program activity can flow to genes, the reference regulatory path narrows the solution space, anchors the model in biologically plausible regulatory structure, and enables mechanistic interpretation of predicted perturbation responses. In contrast, the mapping from perturbation targets to regulatory programs is learned directly from the Perturb-seq data, allowing the model to capture perturbation-specific regulatory effects while reusing a shared regulatory architecture.

REGA-perturb takes four inputs. The first is the normalized gene expression matrix 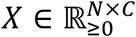, containing *N* genes across *C* cells (see “Data collection and preprocessing”). The second is the perturbation design matrix

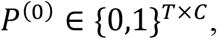

where 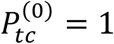 if cell *c* is assigned to perturbation target *t*, and 0 otherwise. Each cell is assigned to exactly one of *T* perturbation categories, including a non-targeting control category.

The remaining two inputs specify the reference regulatory path through RE–motif and RE–TG associations. To identify potential TF binding sites, RE sequences from the reference RE catalog were scanned using HOMER^56^ (findMotifsGenome.pl, with the -find and -size given options; hg38).

The resulting binary RE–motif matrix

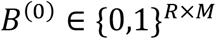

indicates motif occurrence, with 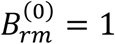 if motif *m* is detected in RE *r*, and 0 otherwise.

To define candidate RE–TG relationships, REs within 300 kb of a gene’s transcription start site (TSS) were considered potential regulators. The initial RE–TG weight matrix 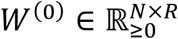 was defined as

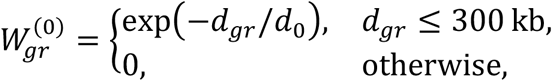

where *d*_*gr*_ denotes the genomic distance between the midpoint of RE *r* and the TSS of gene *g*, and *d*_0_ = 100 kb is the distance-decay parameter. This initialization assigns larger weights to proximal RE–TG pairs.

REGA-perturb models the transcriptional responses to genetic perturbation as a weighted combination of *K* latent regulatory programs:

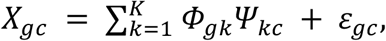

where *Φ*_*gk*_ represents the contribution of program *k* to gene *g, Ψ*_*kc*_ denotes the activity of program *k* in cell *c*, and ε_*gc*_ is a Gaussian error term.

To capture perturbation-specific regulatory activity, each perturbation target *t* is associated with a program activity vector *H*_:,*t*_, where *H* ∈ ℝ ≥ 0^*K×T*^. This activity profile is shared by cells carrying the same perturbation, with *H*_:,ctrl_ representing the profile of non-targeting control cells. To account for cell-to-cell variation in the overall magnitude of program activity, each cell *c* is assigned a positive scaling factor *p*_c_, such that

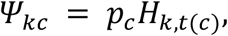

where *t*(*c*) denotes the perturbation category assigned to cell *c*. In matrix form, *Ψ* = *HP*, where *P* ∈ ℝ ≥ 0^*T×C*^ has the same nonzero pattern as the perturbation design matrix *P*^(0)^, with *P*_*tc*_ = *p_c_* if *t* = *t*(*c*) and 0 otherwise.

The contribution of each regulatory program to gene expression is further decomposed through the reference regulatory path. Specifically, *V*_*mk*_ represents the contribution of TF motif *m* to program *k, B*_*rm*_ denotes the weight of motif *m* at RE *r*, and *W*_*gr*_ represents the regulatory contribution of RE *r* to gene *g*. Accordingly,

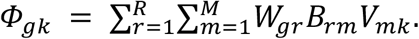

Combining the program-to-gene contributions with perturbation-specific program activities yields a five-factor non-negative matrix factorization^57^:

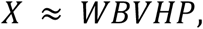

where *W*, *B* and *V* link regulatory programs to genes through TF motifs and REs, *H* represents perturbation-specific program activity profiles, and *P* encodes perturbation assignments with cell-specific scaling factors.

Model parameters were estimated by minimizing the squared reconstruction error with L2 regularization on *W, B, V* and *H*:

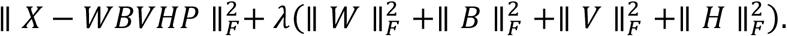

The L2 penalties discourage excessively large factor weights and help stabilize optimization, whereas the sparsity of *W* and *B* is enforced by the reference regulatory path.

We designed an alternating multiplicative update algorithm to estimate the model parameters^58^:

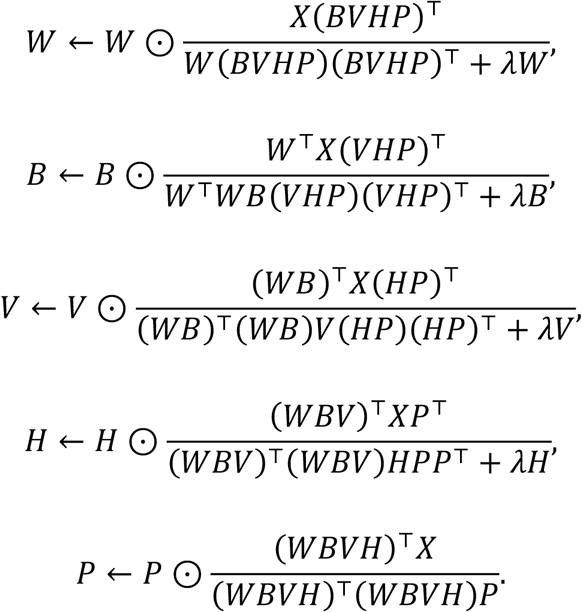

Here, ⊙ and the fractions denote element-wise multiplication and division, respectively.

To accommodate datasets containing millions of cells, we employed mini-batch multiplicative updates. At each epoch, cells were randomly shuffled and partitioned into mini-batches. For each mini-batch ℬ, the parameters were updated sequentially in the order *W, B, V, H* and *P*, using the corresponding subsets *X*_ℬ_ and *P*_ℬ_. The shared factors *W, B* and *V* were updated after each mini-batch, whereas *H* was updated only for perturbation targets represented in the current mini-batch and *P* only for cells in that mini-batch. *W, B* and *P* were initialized with *W*^(0)^, *B*^(0)^ and *P*^(0)^, whereas *V* and *H* were initialized with values drawn from a uniform distribution on [0,1). REGA-perturb was implemented in PyTorch^59^ and trained on GPUs.

### Inference of cis- and trans-regulatory interactions

We represented genes, regulatory elements (REs), transcription factor (TF) motifs, perturbation targets and cells in a shared space of *K* latent regulatory programs. Their embeddings were defined as

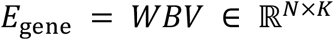

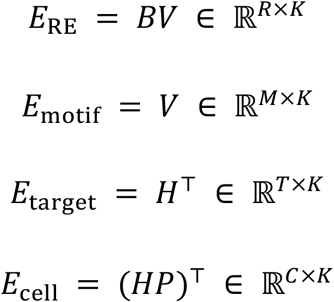

Here, *E*_gene_ summarizes the regulatory program associations of each gene through its linked REs and TF motifs, whereas *E*_RE_ captures the program associations of each RE through its associated motifs. *E*_target_ represents the program activity profile of each perturbation target, and *E*_cell_ incorporates cell-specific scaling factors into these activities. The model reconstructs gene expression as *X* ≈ *E*_gene_*E*_cell_^T^ (see “REGA-perturb model”).

To obtain TF-level embeddings, we mapped motifs to TFs using a binary motif–TF annotation matrix Γ ∈ {0,1}^*M×F*^, where Γ_*mf*_=1 if motif *m* is annotated to TF *f*. Each TF embedding was defined as the sum of its associated motif embeddings:

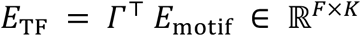

Regulatory relationships were inferred from dot products between these embeddings.

For each candidate RE–target gene pair defined by the reference regulatory path, we calculated an embedding-based regulatory score as

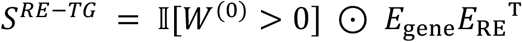

where I[·] denotes the indicator function. This score quantifies the compatibility between gene and RE embeddings based on their shared associations with latent regulatory programs.

We calculated TF–RE association scores from the dot products of their embeddings, restricted to REs containing the corresponding TF motifs:

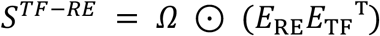

where *Ω*_*rf*_=1 if RE *r* contains at least one motif annotated to TF *f*, as determined by (*B*^(0)^Γ)_*rf*_ > 0, and *Ω*_*rf*_ =0 otherwise. This score captures shared regulatory program associations between TFs and REs supported by motif occurrences.

To infer TF–target gene relationships, we calculated two complementary scores. The functional influence score was defined as

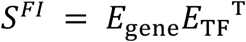

This score captures TF–target gene associations through shared regulatory programs without requiring the presence of the corresponding TF motif in REs linked to the gene.

The regulatory connectivity score instead considers TF–target gene associations supported by candidate regulatory paths through REs:

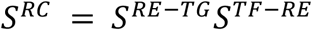

This score integrates RE–target gene regulatory scores and TF–RE association scores, restricting TF– target gene associations to motif-supported regulatory paths.

The regulatory connectivity score was used to benchmark TF–target gene predictions against ChIP-seq data and to distinguish candidate direct from indirect targets of perturbed TFs. The functional influence score was used to predict regulons responsive to unseen TF perturbations and to construct perturbation-specific and differential regulatory networks.

### RE importance

To quantify the importance of individual regulatory elements (REs), we defined an RE importance score that combines their inferred regulatory contributions to target genes with their aggregate activities across perturbation conditions:

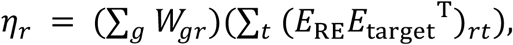

where

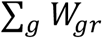

represents the total inferred regulatory weight of RE *r* across target genes, and

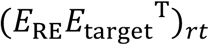

represents its inferred activity under perturbation target *t*, summed over all perturbation targets.

### Perturbation-induced changes in TF activity

The activity of TF *f* under perturbation target *t* was estimated from the dot product of their embeddings, integrating TF loadings across regulatory programs with perturbation-specific program activities:

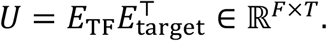

Changes in TF activity relative to non-targeting controls were quantified as log_2_ fold changes:

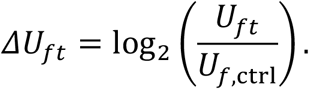

TFs with |*ΔU*_*ft*_| ≥ 0.25 were considered to exhibit altered activity in response to perturbation *t*.

### Perturbation-specific regulatory networks

Within each regulatory program, the scales of *V* and *H* are not identifiable: multiplying *V*_:,*k*_ by a constant and dividing *H*_*k*,:_ by the same constant leaves *WBVHP* unchanged. Before constructing perturbation-specific networks, we therefore rescaled each program so that its motif loadings sum to one, which makes program activities comparable across programs:

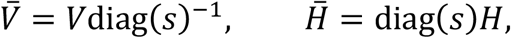

with *s*_*k*_ = ∑_*m*_ *V*_*mk*_. We write 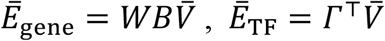, and 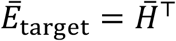 for the corresponding embeddings.

For each perturbation target *t*, we constructed a perturbation-specific TF–TG network by weighting the contribution of each regulatory program according to its activity under perturbation *t*:

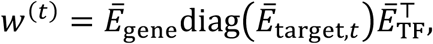

where *Ē*_target,*t*_ denotes the embedding of perturbation target *t*. Equivalently,

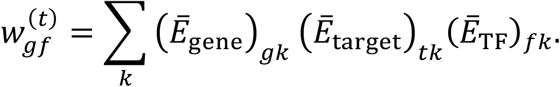

Unlike the functional influence score, which weights all regulatory programs equally, *w*^(*t*)^ incorporates perturbation-specific program activities to quantify TF–TG regulatory associations under each perturbation condition. The corresponding network for non-targeting controls, *w*^(ctrl)^, serves as the reference for evaluating perturbation-associated changes in regulatory influence.

### Perturbation-induced differential networks and rewiring scores

For each perturbation target *t*, we constructed a differential regulatory network by subtracting the network of non-targeting controls from its perturbation-specific network:

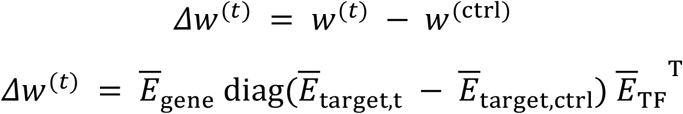

A positive 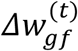 indicates an increase in the inferred regulatory influence of TF *f* on gene *g* under perturbation *t*, whereas a negative value indicates a decrease.

For each gene, we summarized changes in its incoming regulatory interactions across all modeled TFs using two scores: the net regulatory shift score (NRS) and the regulatory rewiring score (RRS):

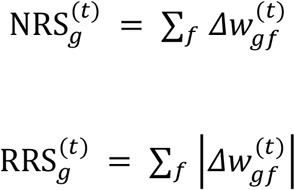

NRS quantifies the net change in inferred regulatory influence, allowing increases and decreases to offset each other, whereas RRS captures the total magnitude of regulatory changes regardless of direction.

### Metacell construction

To make the baseline methods computationally tractable and ensure consistent inputs across methods, we aggregated cells into metacells separately within each perturbation target using SEACells^60^. For benchmarking, both REGA-perturb and the baseline methods were fitted using metacell expression profiles, whereas all other analyses used REGA-perturb models fitted to the complete single-cell datasets.

### Benchmarking trans-regulatory inference

We compared the performance of REGA-perturb in predicting trans-regulatory (TF–TG) interactions with that of PCC, GRNBoost2 (arboreto v0.1.6)^22^, SCENIC (pySCENIC v0.12.1)^20,61^ and CellOracle (v0.20.0)^21^.

### Evaluation metrics

The performance of gene regulatory network inference was evaluated using the area under the precision– recall curve (AUPR), the area under the receiver operating characteristic curve (AUROC), and precision, recall and F1-score based on the top 25% of predicted regulatory interactions. To account for differences in the proportion of positive interactions across benchmark datasets, we also calculated the AUPR ratio, defined as the observed AUPR normalized by the expected AUPR of a random predictor:

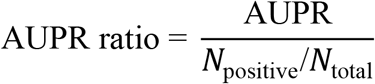

where *N*_positive_ and *N*_total_ denote the numbers of positive and total candidate regulatory interactions, respectively. As the expected AUPR of a random predictor equals the fraction of positive interactions, the AUPR ratio measures the fold enrichment in precision–recall performance relative to random prediction.

## Data availability

All datasets used in this study are publicly available. The K562 genome-wide and essential-gene Perturb-seq datasets from Replogle et al.^3^ were obtained from figshare (https://doi.org/10.25452/figshare.plus.20029387.v1), and the HepG2 Perturb-seq dataset from Nadig et al.^29^ was obtained from the Gene Expression Omnibus (GEO; accession GSE264667). Bulk ATAC-seq data used to construct the reference regulatory element (RE) catalogs were obtained from ENCODE^27^ (https://www.encodeproject.org/) under accession numbers ENCSR956DNB, ENCSR859USB, ENCSR483RKN and ENCSR868FGK for K562, and ENCSR291GJU and ENCSR042AWH for HepG2.

TF ChIP-seq data for both cell lines were also obtained from ENCODE, with accession numbers for all experiments provided in Supplementary Table 1. RNA polymerase II (POLR2A) ChIA-PET data were obtained from ENCODE under accession numbers ENCSR880DSH (K562) and ENCSR857MYZ (HepG2). H3K27ac and H3K4me3 ChIP-seq data for both cell lines were also obtained from ENCODE. Base-wise evolutionary conservation scores (phastCons, 100-way vertebrate alignment; hg38) were downloaded from the UCSC Genome Browser (https://hgdownload.soe.ucsc.edu/goldenPath/hg38/phastCons100way/). TF knockdown and knockout datasets were obtained from KnockTF 2.0^36^ (http://www.licpathway.net/KnockTF/index.html). K562 SNARE-seq data^33^ were obtained from GEO (GSE126074), and human liver single-nucleus multiome data^34^ were obtained from GEO (GSE296875), from which hepatocyte profiles were used in this study.

## Code availability

The software is available at GitHub (https://github.com/Durenlab/REGA-perturb).

## Acknowledgments

This work was partially supported by NIH grant R35GM150513.

## Author Contributions

Z.D. conceived the study. Z.D. and L.R. designed the analytical approach. L.R. performed the data analyses and developed the software. Z.D. supervised the project and acquired funding. L.R. and Z.D. wrote, reviewed and edited the manuscript. Both authors read and approved the final manuscript.

---

**Supplementary Figure 1.**
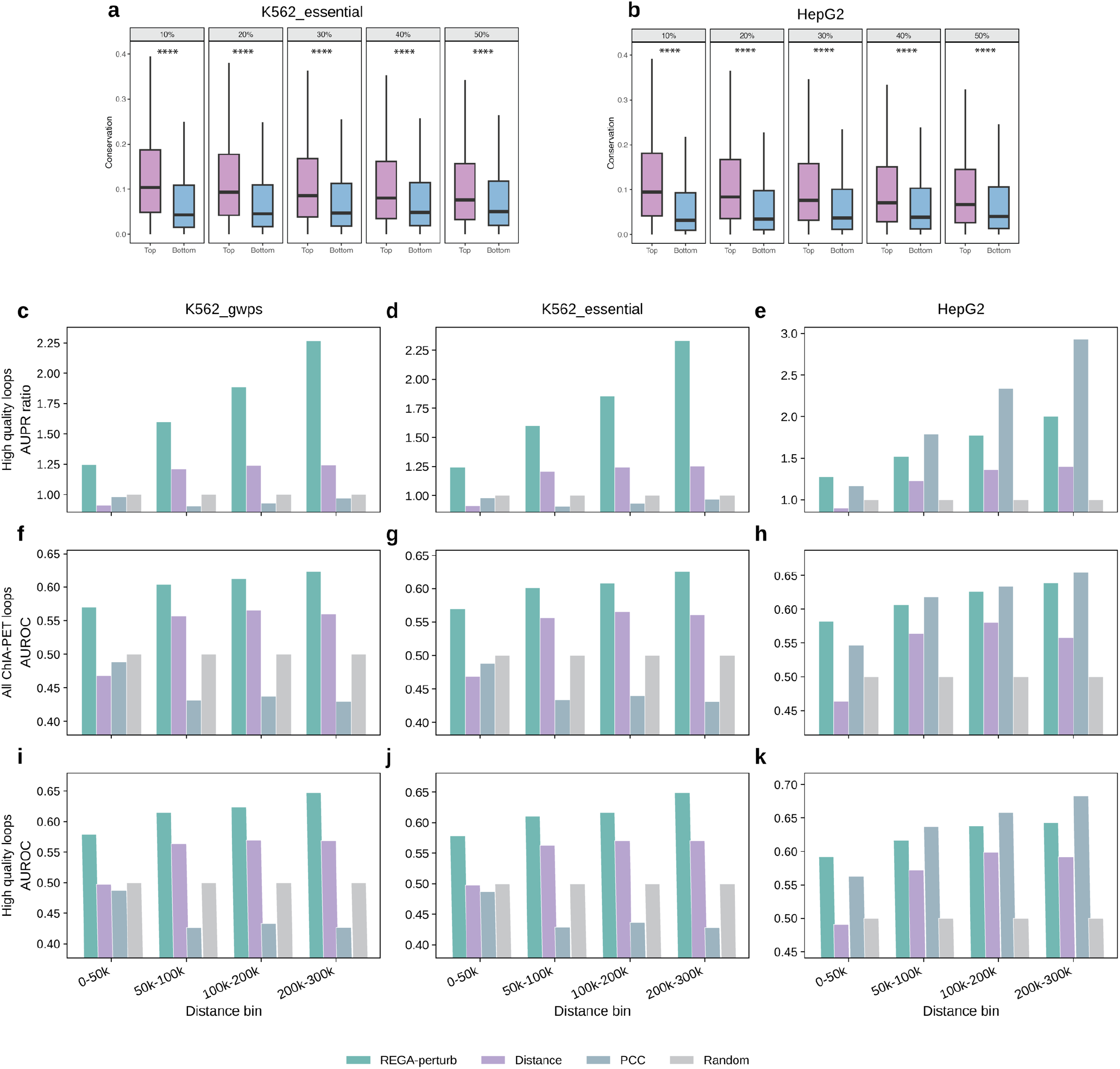
Conservation of prioritized regulatory elements and additional evaluation of RE–target gene inference. **a**,**b**, Sequence conservation of regulatory elements (REs) prioritized by REGA-perturb in the K562 essential-gene screen (K562 essential; **a**) and HepG2 (**b**). Boxplots compare 100-way phastCons conservation scores between REs with high (Top) and low (Bottom) REGA-perturb importance across percentile thresholds of 10%, 20%, 30%, 40% and 50%. Boxes indicate the median and interquartile range (IQR), and whiskers extend to 1.5 × IQR. Boxplot statistics were calculated using REs with conservation scores within the displayed range (0–0.42; approximately 5% of REs excluded), whereas statistical tests included all REs. Outliers are not shown. Statistical significance was assessed using two-sided Welch’s t-tests (**a**, P = 1.2 × 10^−90^, 2.5 × 10^−116^, 3.1 × 10^−116^, 1.3 × 10^−107^ and 7.6 × 10^−88^; **b**, P = 8.6 × 10^−189^, 1.5 × 10^−223^, 5.5 × 10^−216^, 2.0 × 10^−201^ and 3.2 × 10^−168^, respectively; ****P < 0.0001). **c–k**, Performance of REGA-perturb in predicting RE–target gene (RE–TG) interactions across four genomic distance bins (0–50, 50–100, 100–200 and 200–300 kb from the transcription start site), using RNA polymerase II ChIA-PET loops as the benchmark reference. Results are shown for the K562 genome-wide Perturb-seq screen (K562 GWPS; **c**,**f**,**i**), K562 essential (**d**,**g**,**j**) and HepG2 (**e**,**h**,**k**). **c–e**, Area under the precision–recall curve (AUPR) ratios evaluated using high-quality loops. **f–h**, Area under the receiver operating characteristic curve (AUROC) evaluated using all loops. **i–k**, AUROC evaluated using high-quality loops. High-quality loops were defined as those with paired-end tag counts at or above the 75th percentile within each experiment. REGA-perturb was compared with three baselines: Distance (genomic distance between the RE and the transcription start site), PCC (Pearson correlation between RE accessibility and gene expression, calculated using paired single-cell multiome data from K562 cells for both K562 datasets and hepatocytes from human liver tissue for HepG2) and Random (randomly assigned scores). Each bar represents a single metric value calculated across all candidate RE–TG pairs within the corresponding distance bin.

**Supplementary Figure 2.**
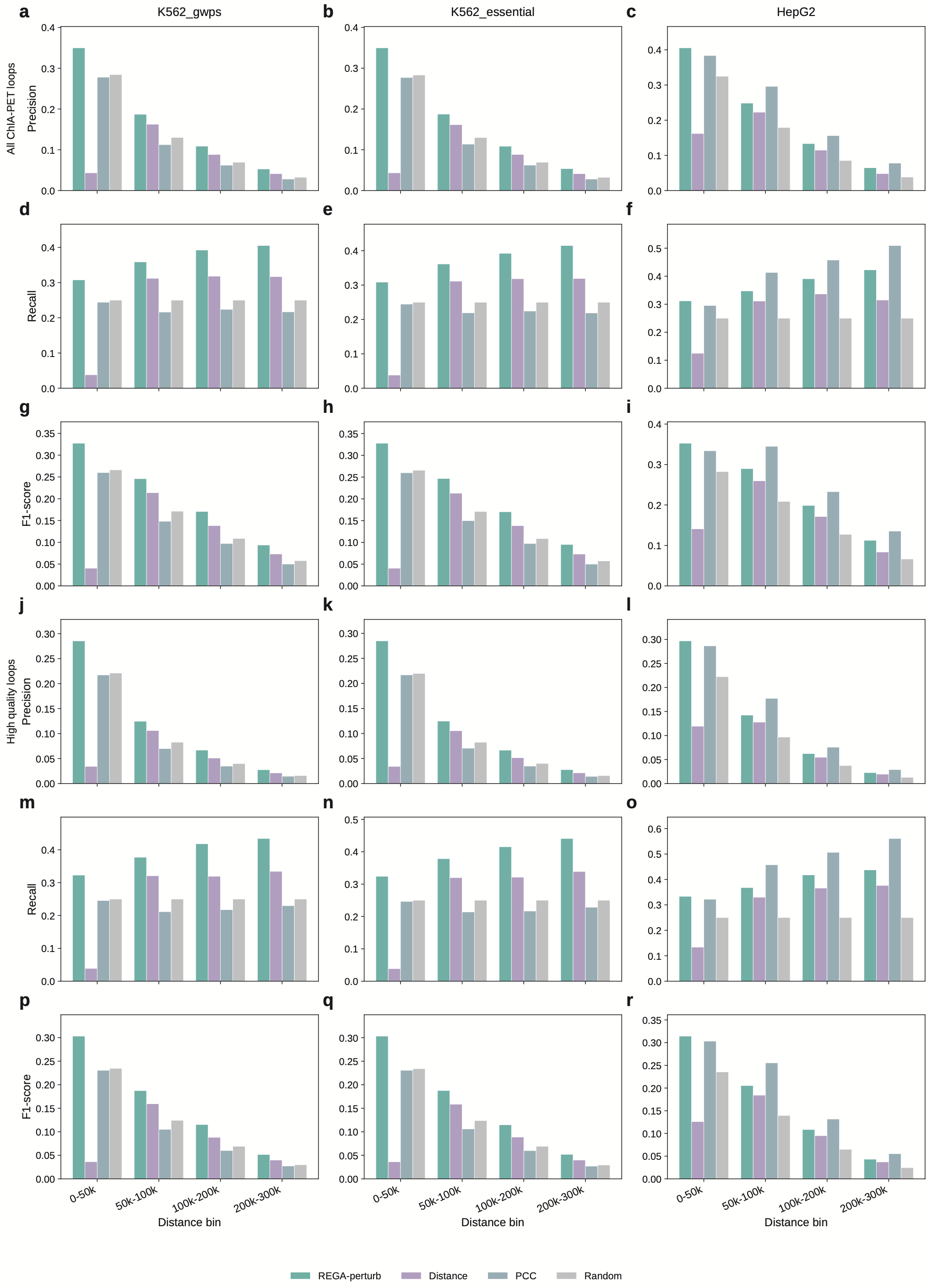
Precision, recall and F1-score of RE–target gene inference by REGA-perturb. Bar plots comparing the performance of REGA-perturb and baseline methods in predicting regulatory element–target gene (RE–TG) interactions across four genomic distance bins (0–50, 50–100, 100–200 and 200–300 kb from the transcription start site), using RNA polymerase II ChIA-PET interactions as the benchmark reference. Results are shown for the K562 genome-wide Perturb-seq screen (K562 GWPS; left column), K562 essential-gene screen (K562 essential; middle column) and HepG2 (right column). **a–i**, Precision (**a–c**), recall (**d–f**) and F1-score (**g–i**) evaluated using all ChIA-PET loops. **j–r**, Corresponding metrics evaluated using high-quality loops: precision (**j–l**), recall (**m–o**) and F1-score (**p–r**). Metrics were calculated using the top 25% of predicted RE–TG interactions ranked within each genomic distance bin. Baseline methods and high-quality loops were defined as in Supplementary Figure 1. Each bar represents a single metric value calculated across all candidate RE–TG pairs within the corresponding distance bin.

**Supplementary Figure 3.**
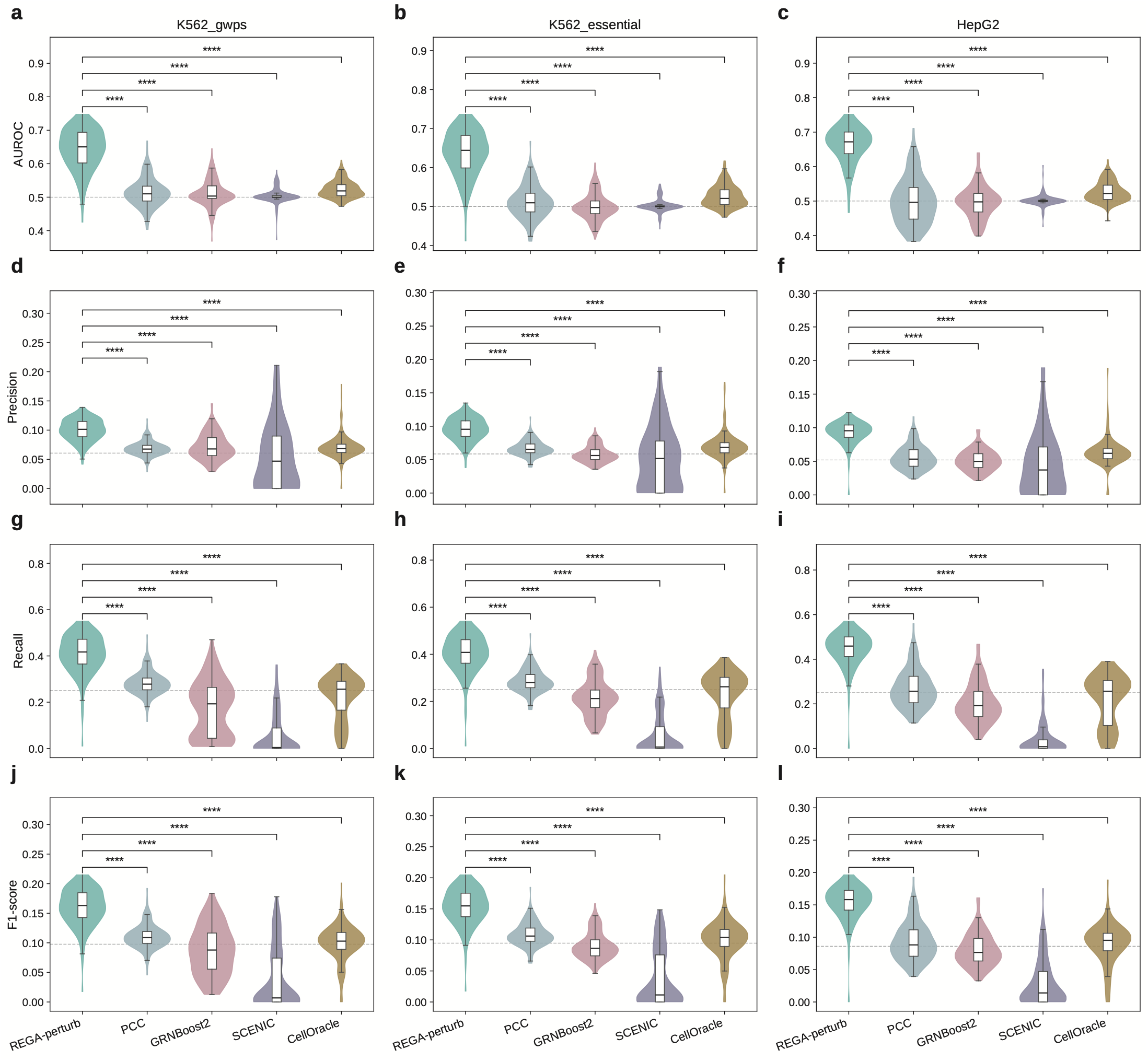
Additional evaluation metrics for TF–target gene inference by REGA-perturb. Violin plots comparing the performance of REGA-perturb, PCC, GRNBoost2, SCENIC and CellOracle in predicting transcription factor–target gene (TF–TG) interactions in the K562 genome-wide Perturb-seq screen (K562 GWPS; **a**,**d**,**g**,**j**), K562 essential-gene screen (K562 essential; **b**,**e**,**h**,**k**) and HepG2 (**c**,**f**,**i**,**l**). ENCODE TF ChIP-seq experiments were used as the benchmark reference. Panels show area under the receiver operating characteristic curve (AUROC; **a–c**), precision (**d–f**), recall (**g–i**) and F1-score (**j–l**). Precision, recall and F1-score were calculated using the top 25% of genes ranked by each method. Boxplots within violins indicate the median and interquartile range (IQR). Dashed lines indicate the expected performance of random rankings (AUROC, 0.5; recall, 0.25; precision, the fraction of positive genes; F1-score, the mean F1-score of random rankings). Violins and boxplots were truncated below the significance brackets for visualization; 1–4 experiments per panel exceeded the displayed range, but all experiments were included in the statistical analyses. Statistical significance was assessed using two-sided paired t-tests across ChIP-seq experiments (n = 152, 157 and 110 experiments from 102, 105 and 88 TFs, respectively; ****P < 0.0001). P values for REGA-perturb versus PCC, GRNBoost2, SCENIC and CellOracle, respectively, were 1.1 × 10^−42^, 3.7 × 10^−49^, 1.6 × 10^−56^ and 2.7 × 10^−51^ in **a**; 1.6 × 10^−48^, 8.4 × 10^−56^, 9.0 × 10^−62^ and 3.9 × 10^−51^ in **b**; 3.0 × 10^−37^, 2.0 × 10^−46^, 7.5 × 10^−54^ and 1.0 × 10^−44^ in **c**; 2.3 × 10^−34^, 1.6 × 10^−23^, 6.5 × 10^−11^ and 2.2 × 10^−25^ in **d**; 2.5 × 10^−38^, 7.0 × 10^−47^, 2.5 × 10^−15^ and 3.0 × 10^−23^ in **e**; 7.1 × 10^−31^, 2.7 × 10^−37^, 2.2 × 10^−14^ and 1.9 × 10^−19^ in **f**; 2.4 × 10^−31^, 2.1 × 10^−44^, 1.7 × 10^−77^ and 2.3 × 10^−44^ in **g**; 5.1 × 10^−34^, 4.5 × 10^−48^, 6.8 × 10^−85^ and 4.0 × 10^−43^ in **h**; 7.1 × 10^−31^, 3.4 × 10^−40^, 5.6 × 10^−66^ and 2.0 × 10^−32^ in **i**; 2.1 × 10^−32^, 3.1 × 10^−37^, 7.4 × 10^−56^ and 2.5 × 10^−38^ in **j**; 1.8 × 10^−35^, 4.3 × 10^−47^, 5.8 × 10^−64^ and 5.4 × 10^−36^ in **k**; and 7.1 × 10^−31^, 3.6 × 10^−39^, 7.8 × 10^−53^ and 4.2 × 10^−29^ in **l**.

**Supplementary Figure 4.**
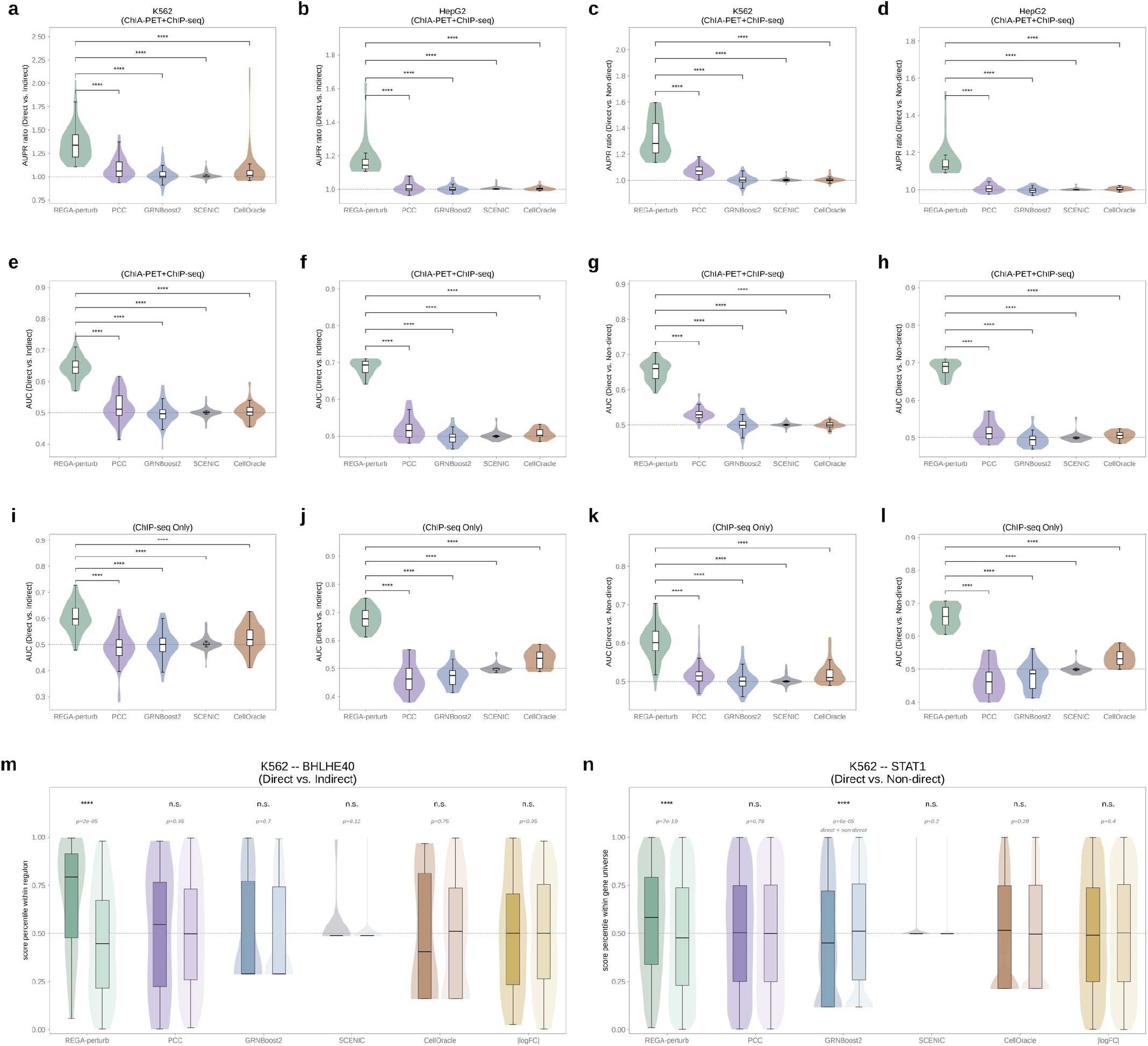
Direct-target identification under alternative target definitions and evaluation metrics. **a–l**, Violin plots comparing the performance of REGA-perturb, PCC, GRNBoost2, SCENIC and CellOracle in distinguishing candidate direct targets of transcription factors (TFs) from other genes. Each violin summarizes TF–ChIP-seq experiment pairs. Columns correspond to the evaluation settings in Figure 3a–d: candidate direct versus indirect targets among differentially expressed genes (DEGs) in K562 (**a**,**e**,**i**) and HepG2 (**b**,**f**,**j**), and candidate direct versus non-direct targets among all modeled genes in K562 (**c**,**g**,**k**) and HepG2 (**d**,**h**,**l**). **a–h**, Candidate direct targets defined using both TF ChIP-seq binding and RNA polymerase II ChIA-PET interactions (ChIA-PET + ChIP-seq), evaluated using area under the precision–recall curve (AUPR) ratios (**a–d**) and area under the receiver operating characteristic curve (AUROC; **e–h**). **i–l**, Candidate direct targets defined using ChIP-seq regulatory potential (RP) scores, as in Figure 3, evaluated using AUROC. Boxplots within violins indicate the median and interquartile range (IQR), and dashed lines indicate random performance (AUPR ratio, 1; AUROC, 0.5). Statistical significance was assessed using two-sided paired Wilcoxon signed-rank tests (K562: n = 67 pairs from 28 TFs, except n = 69 pairs from 29 TFs in **i**; HepG2: n = 20 pairs from 15 TFs; ****P < 0.0001). P values for REGA-perturb versus PCC, GRNBoost2, SCENIC and CellOracle, respectively, were 2.9 × 10^−12^, 1.1 × 10^−12^, 1.1 × 10^−12^ and 5.0 × 10^−11^ in **a**; 1.1 × 10^−12^ for all four comparisons in **c** and **g**; 1.2 × 10^−12^, 1.1 × 10^−12^, 1.1 × 10^−12^ and 1.1 × 10^−12^ in **e**; 9.6 × 10^−12^, 3.4 × 10^−11^, 3.6 × 10^−12^ and 5.5 × 10^−11^ in **i**; and 3.3 × 10^−11^, 5.1 × 10^−12^, 1.9 × 10^−12^ and 4.1 × 10^−12^ in **k**. All 24 comparisons across the six HepG2 panels (**b**,**d**,**f**,**h**,**j**,**l**) yielded P = 1.9 × 10^−6^, the minimum attainable two-sided exact P value for 20 nonzero pairs, with REGA-perturb outperforming each baseline in every paired experiment. **m**,**n**, Additional representative TFs, displayed as in Figure 3e,f. **m**, BHLHE40 in K562 GWPS, comparing candidate direct and indirect targets among Perturb-seq DEGs (P = 1.7 × 10^−5^ for REGA-perturb, 0.95 for PCC, 0.70 for GRNBoost2, 0.12 for SCENIC, 0.75 for CellOracle and 0.95 for absolute log fold change (|logFC|)). **n**, STAT1 in K562 GWPS, comparing candidate direct and non-direct targets among all modeled genes (P = 7.0 × 10^−19^, 0.76, 5.9 × 10^−5^, 0.20, 0.28 and 0.40, respectively). Although GRNBoost2 showed a significant difference, its scores were higher for non-direct than for candidate direct targets. P values were calculated using two-sided Mann–Whitney U tests (n.s., P ≥ 0.05; ****P < 0.0001).

**Supplementary Figure 5.**
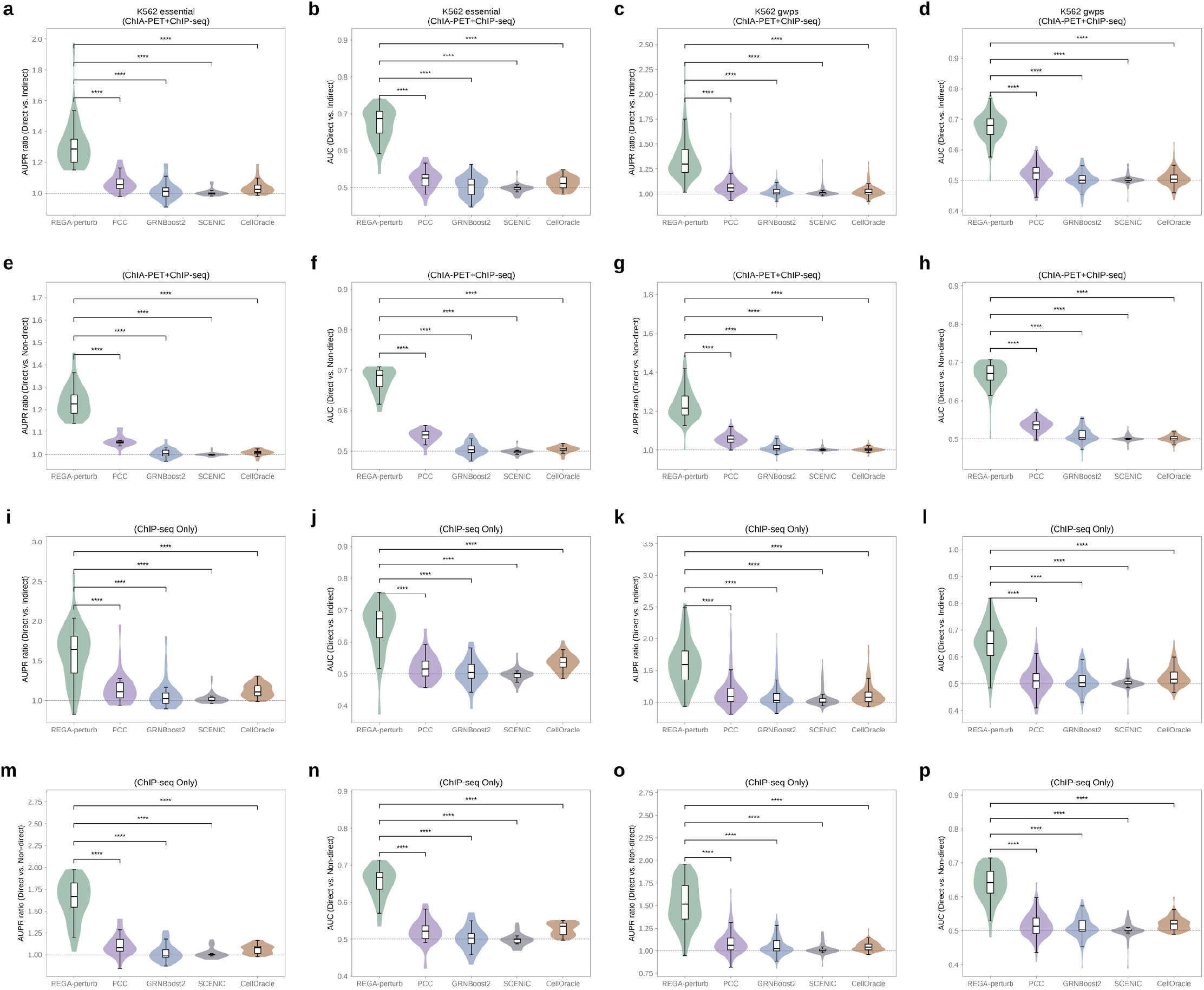
Direct-target identification in K562 using differentially expressed genes from matched Perturb-seq screens. Violin plots comparing the performance of REGA-perturb, PCC, GRNBoost2, SCENIC and CellOracle in distinguishing candidate direct targets of transcription factors (TFs) from other genes in K562, using differentially expressed genes (DEGs) derived from pseudobulk Perturb-seq data rather than the independent KnockTF experiments used in Figure 3a and Supplementary Figure 4. Results are shown for the K562 essential-gene screen (K562 essential; first two columns) and genome-wide screen (K562 GWPS; last two columns), evaluated using area under the precision–recall curve (AUPR) ratios (first and third columns) and area under the receiver operating characteristic curve (AUROC; second and fourth columns). Candidate direct targets were defined using TF ChIP-seq binding supported by RNA polymerase II ChIA-PET interactions (ChIA-PET + ChIP-seq; **a–h**) or ChIP-seq regulatory potential (RP) scores alone (ChIP-seq only; **i–p**). Panels compare candidate direct and indirect targets among DEGs (**a–d**,**i–l**) and candidate direct and non-direct targets among all modeled genes (**e–h**,**m–p**). Each violin summarizes TF–ChIP-seq experiment pairs (K562 essential, n = 27–36 pairs from 17–23 TFs; K562 GWPS, n = 111–151 pairs from 79–101 TFs). Boxplots within violins indicate the median and interquartile range (IQR), and dashed lines indicate random performance (AUPR ratio, 1; AUROC, 0.5). Statistical significance was assessed using two-sided paired Wilcoxon signed-rank tests (****P < 0.0001). The numbers of TF– ChIP-seq experiment pairs (TFs) and P values for REGA-perturb versus PCC, GRNBoost2, SCENIC and CellOracle, respectively, were as follows: **a**,**b**, 35 (22), P = 5.8 × 10^−11^ for all four comparisons; **c**, 149 (99), P = 1.7 × 10^−25^, 3.5 × 10^−26^, 3.6 × 10^−26^ and 3.6 × 10^−26^; **d**, 149 (99), P = 6.0 × 10^−26^, 3.6 × 10^−26^, 3.5 × 10^−26^ and 3.4 × 10^−26^; **e**,**f**, 27 (17), P = 1.5 × 10^−8^ for all four comparisons; **g**, 111 (79), P = 6.3 × 10^−20^, 6.2 × 10^−20^, 6.0 × 10^−20^ and 6.2 × 10^−20^; **h**, 111 (79), P = 6.2 × 10^−20^, 6.2 × 10^−20^, 6.0 × 10^−20^ and 6.2 × 10^−20^; **i**, 36 (23), P = 3.8 × 10^−7^, 5.9 × 10^−8^, 9.6 × 10^−10^ and 6.0 × 10^−9^; **j**, 36 (23), P = 8.9 × 10^−9^, 1.6 × 10^−9^, 5.5 × 10^−10^ and 3.7 × 10^−8^; **k**, 151 (101), P = 2.7 × 10^−20^, 6.3 × 10^−22^, 1.7 × 10^−24^ and 3.3 × 10^−23^; **l**, 151 (101), P = 9.9 × 10^−24^, 2.9 × 10^−24^, 1.7 × 10^−25^ and 5.5 × 10^−24^; **m**, 27 (17), P = 1.5 × 10^−8^, 4.5 × 10^−8^, 3.0 × 10^−8^ and 3.0 × 10^−8^; **n**, 27 (17), P = 1.5 × 10^−8^ for all four comparisons; **o**, 111 (79), P = 3.3 × 10^−18^, 1.9 × 10^−19^, 9.2 × 10^−20^ and 1.0 × 10^−19^; **p**, 111 (79), P = 6.4 × 10^−19^, 9.2 × 10^−20^, 7.0 × 10^−20^ and 8.1 × 10^−20^.

**Supplementary Figure 6.**
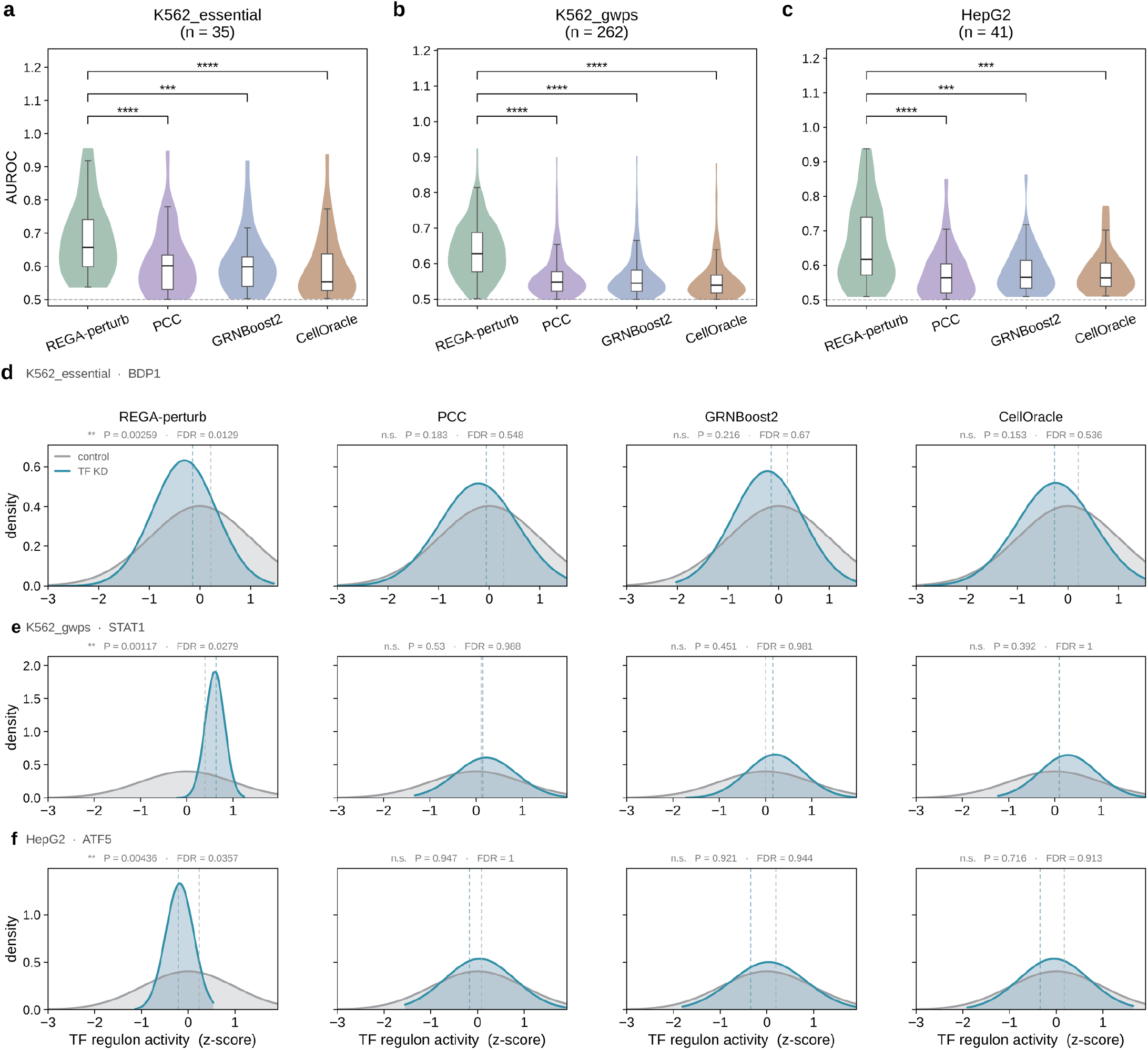
AUROC evaluation and additional examples for unseen TF perturbations. **a–c**, Violin plots comparing the area under the receiver operating characteristic curve (AUROC) for distinguishing knockdown from control cells based on regulon activity for transcription factors (TFs) whose perturbations were excluded from model training (unseen TFs). Results are shown for the K562 essential-gene screen (K562 essential; **a**, n = 35 unseen TFs), K562 genome-wide screen (K562 GWPS; **b**, n = 262) and HepG2 (**c**, n = 41). Regulon definitions, activity calculations and selection of ranking direction were as described in Figure 4. Paired comparisons used the unseen TFs shared by REGA-perturb and each baseline (n = 34, 249 and 37 for CellOracle in **a, b** and **c**, respectively). Boxplots within violins indicate the median and interquartile range (IQR), and the dashed line indicates an AUROC of 0.5. Mean AUROC values for REGA-perturb, PCC and GRNBoost2 were 0.68, 0.61 and 0.61 in **a**; 0.64, 0.56 and 0.56 in **b**; and 0.66, 0.58 and 0.59 in **c**, respectively. Corresponding values for CellOracle were 0.60, 0.56 and 0.58. Statistical significance was assessed using two-sided paired t-tests across unseen TFs (REGA-perturb versus PCC, GRNBoost2 and CellOracle: P = 1.2 × 10^−5^, 1.8 × 10^−4^ and 6.1 × 10^−6^ in **a**; 1.1 × 10^−42^, 1.1 × 10^−42^ and 1.3 × 10^−45^ in **b**; and 2.6 × 10^−5^, 3.3 × 10^−4^ and 1.3 × 10^−4^ in **c**; ***P < 0.001, ****P < 0.0001). **d–f**, Additional examples of unseen TF perturbations: BDP1 in K562 essential (**d**), STAT1 in K562 GWPS (**e**) and ATF5 in HepG2 (**f**). Density plots show regulon activity, expressed as z-scores relative to controls, in knockdown and control cells for regulons predicted by REGA-perturb, PCC, GRNBoost2 and CellOracle (left to right). Curves represent Gaussian fits, and dashed lines indicate group medians. P values were calculated using two-sided Mann–Whitney U tests, with false discovery rates (FDRs) estimated using the Benjamini–Hochberg procedure across all unseen TFs evaluated by each method. For REGA-perturb, PCC, GRNBoost2 and CellOracle, respectively, P values (FDRs) were 2.6 × 10^−3^ (0.013), 0.18 (0.55), 0.22 (0.67) and 0.15 (0.54) in **d**; 1.2 × 10^−3^ (0.028), 0.53 (0.99), 0.45 (0.98) and 0.39 (1.00) in **e**; and 4.4 × 10^−3^ (0.036), 0.95 (1.00), 0.92 (0.94) and 0.72 (0.91) in **f** (n.s., P ≥ 0.05; **P < 0.01).

**Supplementary Figure 7.**
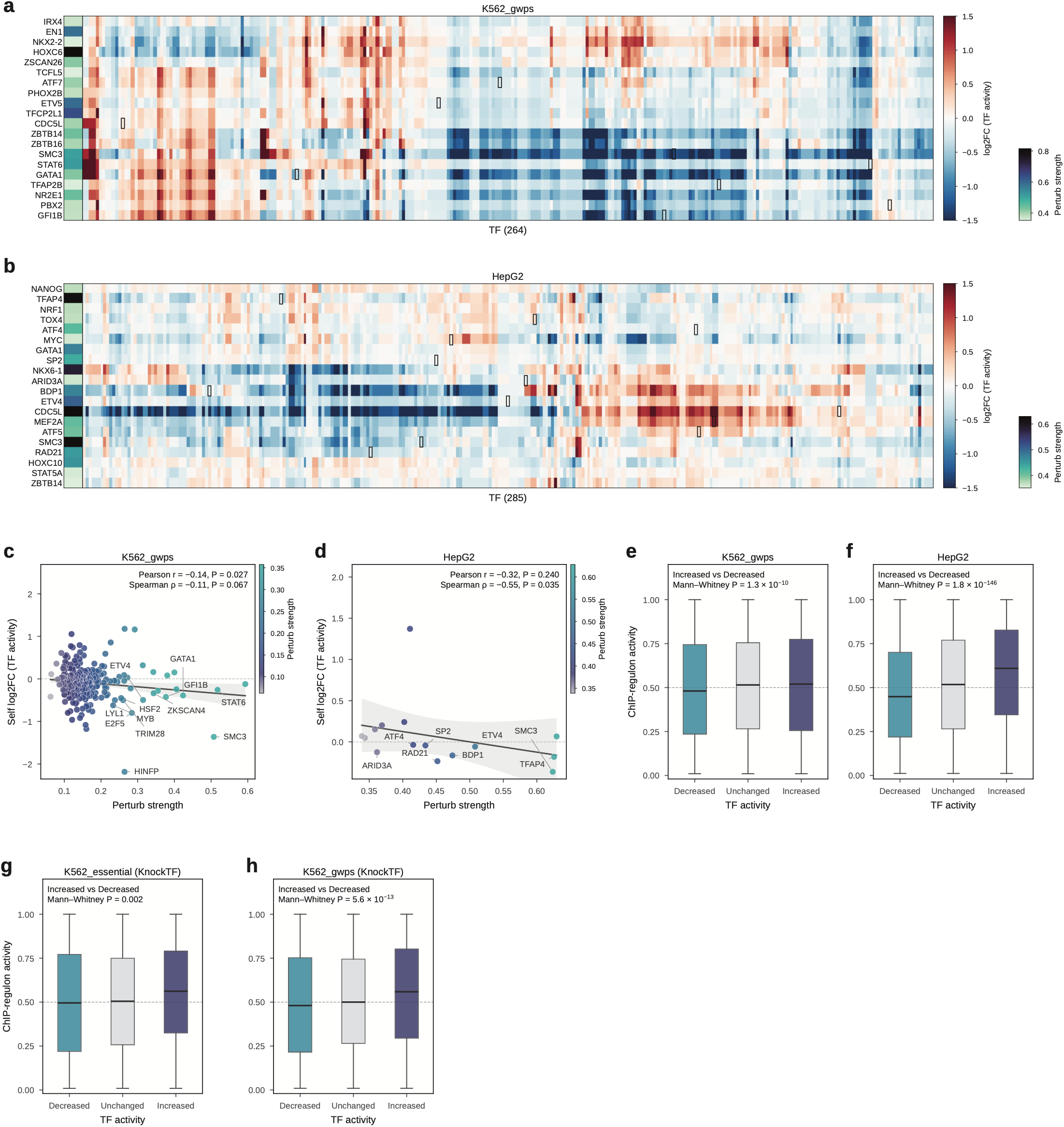
Perturbation-induced changes in TF activity across additional datasets and independent knockdown experiments. **a**,**b**, Heatmaps showing log_2_ fold changes (log_2_FCs) in REGA-perturb-inferred transcription factor (TF) activity relative to non-targeting controls following knockdown of the 20 TFs with the strongest perturbation effects (rows) in the K562 genome-wide screen (K562 GWPS; **a**, 264 modeled TFs) and HepG2 (**b**, 285 modeled TFs), displayed as in Figure 5a. Black boxes indicate the perturbed TF itself in 9 and 13 of the 20 rows in **a** and **b**, respectively; the remaining perturbed TFs were not included in the model. **c**,**d**, Scatter plots comparing changes in the perturbed TF’s own inferred activity with perturbation strength for 264 TF perturbations in K562 GWPS (**c**) and 15 TF perturbations within the top 30% of perturbation strength in HepG2 (**d**), displayed as in Figure 5b (two-sided correlation tests; **c**, Pearson r = −0.14, P = 0.027; Spearman ρ = −0.11, P = 0.067; **d**, Pearson r = −0.32, P = 0.24; Spearman ρ = −0.55, P = 0.035). **e**,**f**, Boxplots comparing ChIP-regulon activity among TFs predicted to have decreased (log_2_FC ≤ −0.25), unchanged (|log_2_FC| < 0.25) or increased (log_2_FC ≥ 0.25) activity across the 500 strongest perturbations in K562 GWPS (**e**; n = 16,873, 25,498 and 8,624 knockdown–TF pairs, respectively) and HepG2 (**f**; n = 16,115, 21,804 and 5,577), using ChIP-regulon activity calculated from the corresponding Perturb-seq datasets as in Figure 5c. **g**,**h**, Corresponding analyses using ChIP-regulon activity derived from independent bulk knockdown experiments in the KnockTF database, evaluated with predictions from the K562 essential-gene model (**g**; 66 experiments; n = 832, 5,585 and 495 knockdown–TF pairs) and K562 GWPS model (**h**; 180 experiments; n = 4,547, 10,810 and 2,955). Boxplots are displayed as in Figure 5c. Median ChIP-regulon activities for TFs with decreased, unchanged and increased predicted activity were 0.48, 0.51 and 0.52 in **e**; 0.45, 0.52 and 0.61 in **f**; 0.50, 0.50 and 0.56 in **g**; and 0.48, 0.50 and 0.56 in **h**. Statistical significance was assessed using two-sided Mann–Whitney U tests comparing TFs with increased and decreased predicted activity (P = 1.3 × 10^−10^ in **e**, 1.8 × 10^−146^ in **f**, 0.002 in **g** and 5.6 × 10^−13^ in **h**).

**Supplementary Figure 8.**
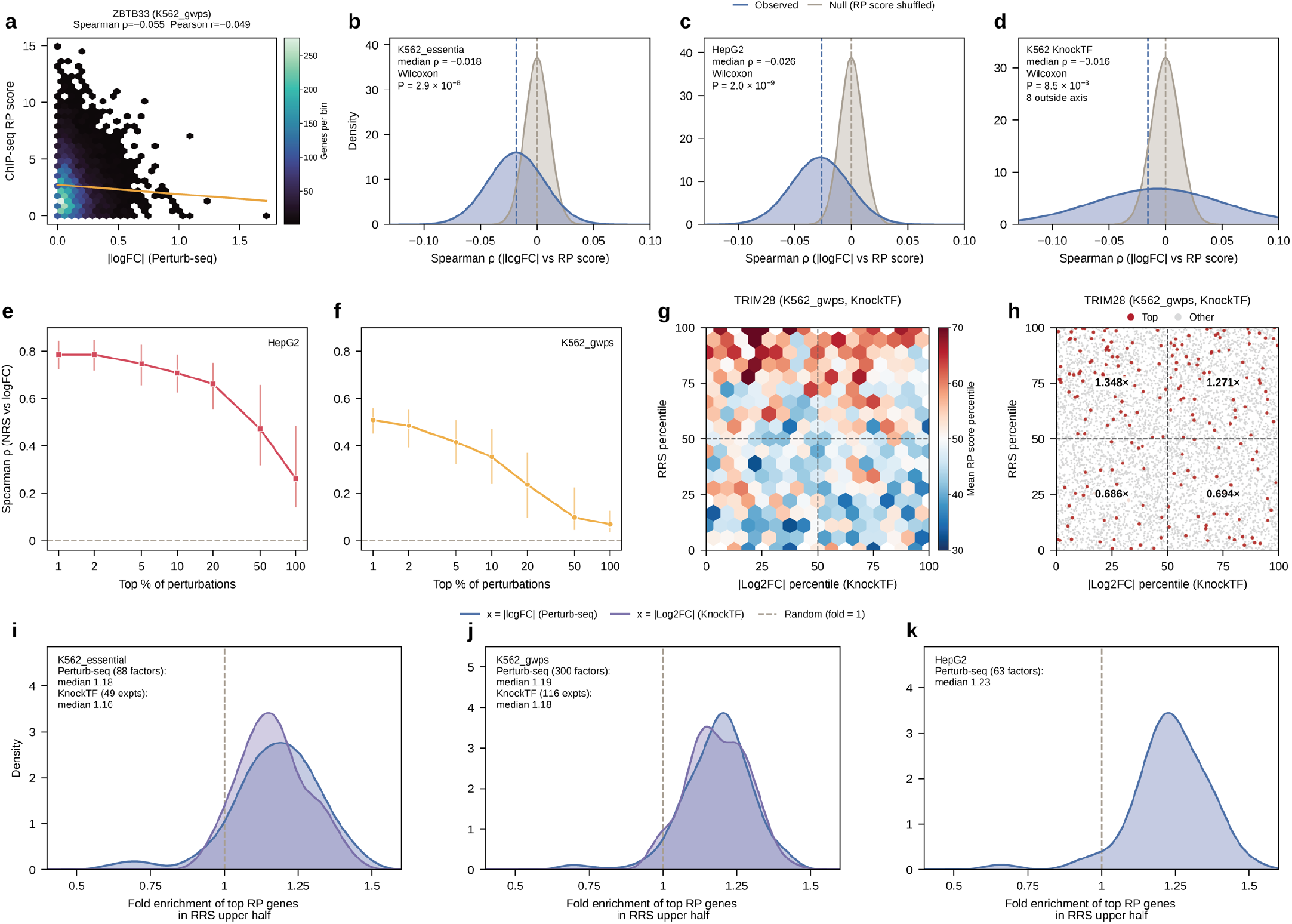
Associations of expression changes, net regulatory shifts and regulatory rewiring with TF binding across additional datasets. **a**, Hexbin plot showing the relationship between the ChIP-seq regulatory potential (RP) scores of ZBTB33 and absolute gene expression changes (|logFC|) following ZBTB33 knockdown in the K562 genome-wide Perturb-seq screen (K562 GWPS; n = 8,227 genes; Spearman ρ = −0.055, two-sided P = 5.9 × 10^−7^). **b–d**, Distributions of Spearman correlations between absolute expression changes (|logFC|) following knockdown and the RP scores of the perturbed transcription factors (TFs) across genes, compared with correlations obtained by shuffling RP scores across genes, in the K562 essential-gene screen (K562 essential; **b**, n = 88 TFs), HepG2 (**c**, n = 63 TFs) and independent bulk knockdown experiments from KnockTF in K562 (**d**, n = 118 experiments targeting 96 TFs). Blue indicates observed correlations, gray indicates shuffled correlations, and dashed lines indicate medians (two-sided paired Wilcoxon signed-rank tests, P = 2.9 × 10^−8^ in **b**, 2.0 × 10^−9^ in **c** and 8.5 × 10^−3^ in **d**). **e**,**f**, Spearman correlations between the net regulatory shift score (NRS) and gene expression changes (logFC) across genes for increasing proportions (top 1–100%) of perturbations ranked by their inferred regulatory changes, as in Figure 6e, in HepG2 (**e**; n = 23–2,343 perturbations) and K562 GWPS (**f**; n = 98–9,849). Points indicate medians, and vertical lines indicate interquartile ranges (IQRs). **g**,**h**, Comparison of absolute gene expression changes following TRIM28 knockdown in an independent bulk KnockTF experiment with regulatory rewiring scores (RRSs) inferred from the K562 GWPS model. Genes are positioned by percentile ranks of |log_2_FC| (x axis) and RRS (y axis). **g**, Hexbin plot colored by the mean percentile rank of TRIM28 ChIP-seq RP scores within each bin. **h**, Scatter plot highlighting the 200 genes with the highest TRIM28 RP scores in red, with fold enrichment indicated for each quadrant (one-sided binomial test, n = 200 genes; P = 3.4 × 10^−3^ and 0.019 for the high-RRS, low-|log_2_FC| and high-RRS, high-|log_2_FC| quadrants, respectively). **i–k**, Distributions of fold enrichment of the top 200 genes ranked by the perturbed TF’s RP scores among genes in the upper half of RRS, across TF perturbations in K562 essential (**i**), K562 GWPS (**j**) and HepG2 (**k**). Blue indicates TF perturbations evaluated using the corresponding Perturb-seq datasets (n = 88, 300 and 63, respectively), and purple indicates independent KnockTF experiments evaluated using the K562 essential and K562 GWPS models (n = 49 and 116, respectively). The dashed line indicates a fold enrichment of 1. Statistical significance was assessed using two-sided Wilcoxon signed-rank tests against 1 (Perturb-seq: P = 1.8 × 10^−12^, 1.6 × 10^−44^ and 6.8 × 10^−11^ in **i–k**, respectively; KnockTF: P = 2.9 × 10^−9^ and 3.6 × 10^−20^ in **i**,**j**, respectively).

**Supplementary Figure 9.**
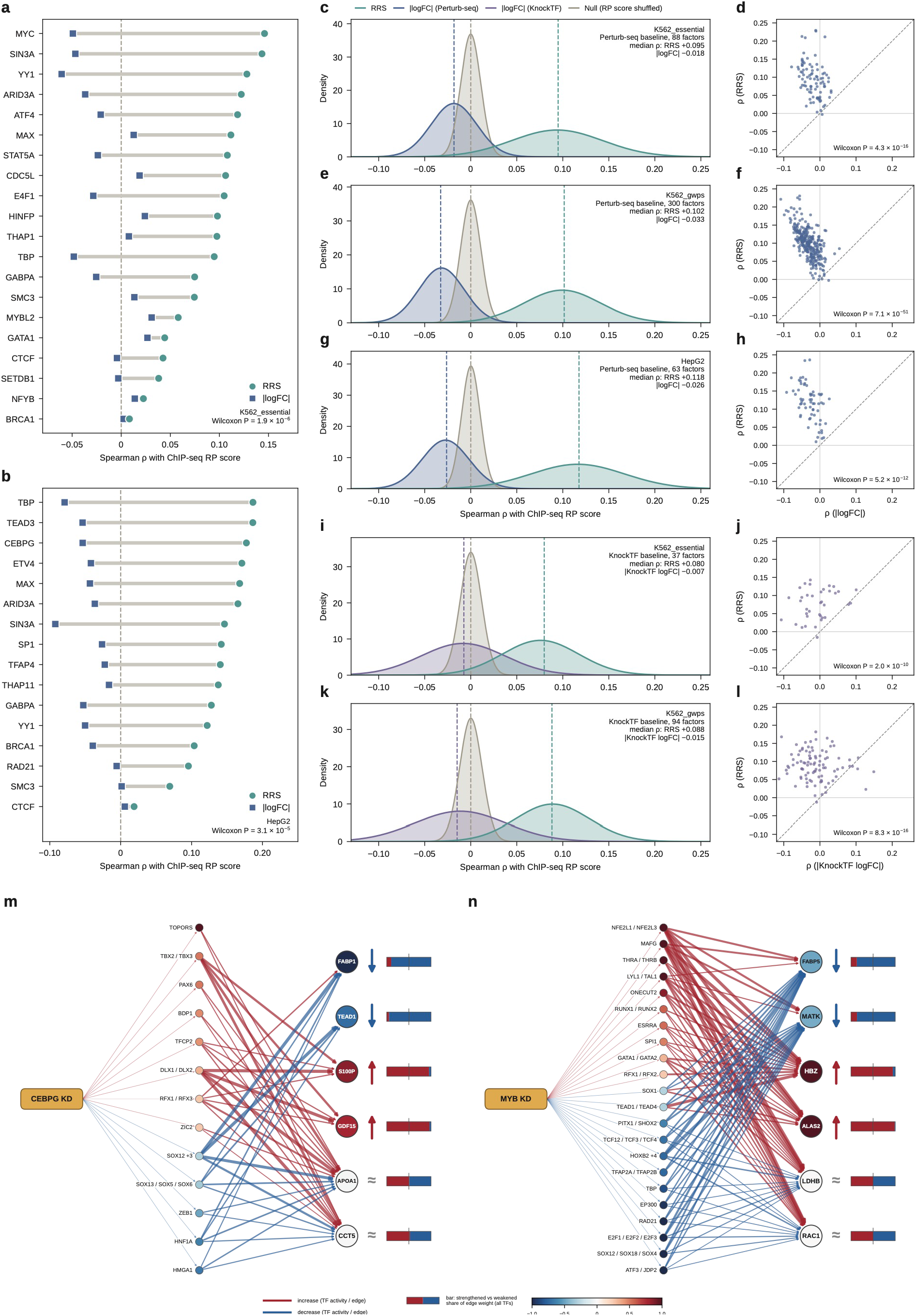
Regulatory rewiring scores are more strongly associated with TF binding than expression changes. **a**,**b**, Dumbbell plots comparing Spearman correlations of regulatory rewiring scores (RRSs; circles) and absolute expression changes (|logFC|; squares) with the ChIP-seq regulatory potential (RP) scores of perturbed transcription factors (TFs) across genes in the K562 essential-gene screen (K562 essential; **a**, n = 20 TFs) and HepG2 (**b**, n = 16 TFs). TFs were selected as in Figure 6h, with all 16 eligible TFs included for HepG2 (two-sided paired Wilcoxon signed-rank tests, P = 1.9 × 10^−6^ in **a** and 3.1 × 10^−5^ in **b**). **c–l**, Comparison of RRS and expression changes in their associations with TF binding across all eligible TF perturbations. Density plots (**c**,**e**,**g**,**i**,**k**) show distributions of Spearman correlations with the perturbed TF’s RP scores across genes for RRS (teal), absolute expression changes from the corresponding Perturb-seq screen (blue, |logFC|), independent bulk knockdown experiments from KnockTF (purple, |log_2_FC|) and shuffled RP scores (gray). Curves represent normal fits, and dashed lines indicate medians. Scatter plots (**d**,**f**,**h**,**j**,**l**) compare the correlations of RRS (y axis) and absolute expression changes (x axis) for individual TFs, with the dashed line indicating y = x. Results are shown for K562 essential (**c**,**d**; n = 88 TFs), K562 GWPS (**e**,**f**; n = 300) and HepG2 (**g**,**h**; n = 63) using Perturb-seq expression changes, and for the K562 essential model (**i**,**j**; n = 37) and K562 GWPS model (**k**,**l**; n = 94) using independent KnockTF expression changes. Statistical significance was assessed using two-sided paired Wilcoxon signed-rank tests comparing RRS with expression changes across TFs (P = 4.3 × 10^−16^ in **d**, 7.1 × 10^−51^ in **f**, 5.2 × 10^−12^ in **h**, 2.0 × 10^−10^ in **j** and 8.3 × 10^−16^ in **l**). **m**,**n**, Differential regulatory networks for six target genes following CEBPG knockdown in HepG2 (**m**) and MYB knockdown in K562 GWPS (**n**), visualized as in Figure 6i. Target genes are arranged by decreased, increased and nearly unchanged expression from top to bottom: FABP1, TEAD1, S100P, GDF15, APOA1 and CCT5 in **m**; FABP5, MATK, HBZ, ALAS2, LDHB and RAC1 in **n**. The TF activity color scale is limited to ±1.

